# Multi-omic Analysis of Developing Human Retina and Organoids Reveals Cell-Specific Cis-Regulatory Elements and Mechanisms of Non-Coding Genetic Disease Risk

**DOI:** 10.1101/2021.07.31.454254

**Authors:** Eric D. Thomas, Andrew E. Timms, Sarah Giles, Sarah Harkins-Perry, Pin Lyu, Thanh Hoang, Jiang Qian, Victoria Jackson, Melanie Bahlo, Seth Blackshaw, Martin Friedlander, Kevin Eade, Timothy J. Cherry

**Affiliations:** Center for Developmental Biology and Regenerative Medicine, Seattle Children’s Research Institute, Seattle, WA 98101, USA; Lowy Medical Research Institute, La Jolla, CA 92037, USA; The Scripps Research Institute, La Jolla, CA, 92037, USA; Solomon H. Snyder Department of Neuroscience, Johns Hopkins University School of Medicine, Baltimore, MD 21205, USA; Department of Ophthalmology, Wilmer Eye Institute Johns Hopkins University School of Medicine, Baltimore, MD 21205, USA; Walter and Eliza Hall Institute, Melbourne, Australia; Kavli Neuroscience Discovery Institute, Johns Hopkins University, Baltimore, MD 21218, USA; Department of Neurology, Johns Hopkins University School of Medicine, Baltimore, MD 21205, USA; Center for Human Systems Biology, Johns Hopkins University School of Medicine, Baltimore, MD 21205, USA; Department of Pediatrics, University of Washington School of Medicine, Seattle, WA 98195, USA; Department of Biological Structure, University of Washington School of Medicine, Seattle, WA 98195, USA; Department of Ophthalmology, University of Washington School of Medicine, Seattle, WA 98195, USA; Brotman Baty Institute, Seattle, WA 98195, USA

**Keywords:** Retina, Retinal Organoid, Development, Cis-regulatory element, MIR-9, Macular Telangiectasia Type 2, Neurogenesis, Single Cell ATAC-seq, Single Cell RNA-seq

## Abstract

Cis-regulatory elements (CREs) play a critical role in the development, maintenance, and disease-states of all human cell types. In the human retina, CREs have been implicated in a variety of inherited retinal disorders. To characterize cell-class-specific CREs in the human retina and elucidate their potential functions in development and disease, we performed single-nucleus (sn)ATAC-seq and snRNA-seq on the developing and adult human retina and on human retinal organoids. These analyses allowed us to identify cell-class-specific CREs, enriched transcription factor binding motifs, putative target genes, and to examine how these features change over development. By comparing DNA accessibility between the human retina and retinal organoids we found that CREs in organoids are highly correlated at the single-cell level, validating the use of organoids as a model for studying disease-associated CREs. As a proof of concept, we studied the function of a disease-associated CRE at 5q14.3 in organoids, identifying its principal target gene as the miR-9-2 primary transcript and demonstrating a dual role for this CRE in regulating neurogenesis and gene regulatory programs in mature glia. This study provides a rich resource for characterizing cell-class-specific CREs in the human retina and showcases retinal organoids as a model in which to study the function of retinal CREs that influence retinal development and disease.

**HIGHLIGHTS:**

1. Single-cell map of cis-regulatory elements in developing and adult human retina.
2. Correlation of single-cell DNA accessibility between human retina and retinal organoids.
3. Association of disease risk loci with cell-class-specific accessibility.
4. Modeling of enhancer function at the 5q14.3 retinal disease-risk locus.

## INTRODUCTION

Understanding the role of the non-coding genome in human development and disease is a persistent challenge due to its scope and complexity. CREs, including enhancers, promoters, silencers, and boundary elements, are essential for controlling the dynamic programs of gene expression necessary for the specification and mature function of distinct cell types. Genetic variants that disrupt CRE function have been associated with a variety of human diseases (Spielmann and Mundlos, 2016). However, the genomic location and function of the majority of CREs is still unclear, largely because CREs can be cell-type-specific, developmentally dynamic, located at a great distance from their target genes, and often are not conserved across model systems. Genome-wide assays now make it possible to identify candidate CREs based on their epigenomic signatures including hyper-accessibility of DNA to nucleases or transposases (Buenrostro et al., 2013; Thurman et al., 2012). As single-cell sequencing technologies have advanced, it is now possible to examine programs of cell-type-specific gene expression in a vast number of diverse cell types and also identify cell-type-specific patterns of chromatin accessibility, which will aid in the study of CREs in development, homeostasis, and disease.

The retina has long served as a classic model system in which to study the development and temporal order of specification of a diverse array of neural cell classes. The gene regulatory networks (GRNs) that control the specification of retinal cell classes from multipotent progenitors represent an area of active research (Andzelm et al., 2015; Chan et al., 2020; Lyu, 2021). More recently, the role of CREs in this process has become an important focus (Aldiri et al., 2017; Chan et al., 2020; Goodson et al., 2020; Lyu, 2021; Norrie et al., 2019; Wang et al., 2014). Furthermore, several studies have implicated mutations in regulatory regions in developmental and mature-onset retinal disorders such as blue cone monochromacy, aniridia with foveal hypoplasia, non-syndromic retinal non-attachment, and age-related macular degeneration (AMD) (Bhatia et al., 2013; Ghiasvand et al., 2011; Han et al., 2020; Liao et al., 2017; Nathans et al., 1989). However, determining the functional role of human retinal CREs within their native chromatin context and in human retinal-like cells has been limited by the availability of validated experimental models.

Retinal organoids derived from human induced pluripotent stem cells (hiPSCs) have emerged as a promising model for studying human retinal development. Organoids recapitulate key aspects of the cytoarchitecture, cellular diversity, and developmental gene expression programs of the human retina (Capowski et al., 2019; Eiraku et al., 2011; Lu et al., 2020; Sridhar et al., 2020). A recent notable study compared the global DNA accessibility landscape of developing organoids to the developing human retina and found broad similarities, but this dataset lacked single-cell resolution as well as comparison to adult retina (Xie et al., 2020). If retinal cell-class-specific CRE activity is recapitulated by retinal organoids, then they could serve as an excellent model system for functional studies of human retinal CREs.

In this study, we sought to identify cell-class-specific CREs in the developing and mature human retina by performing snATAC-seq and comparing these tissues to developing human retinal organoids to benchmark this model for studies of human retinal CREs. We coupled these studies with snRNA-seq analysis to aid in the identification of retinal cell classes, cell-class-specific regions of accessibility, and enriched transcription factor binding motifs, as well as to identify putative target genes for individual CREs. Correlation of our human and organoid datasets demonstrates that the cell-class-specific DNA accessibility landscape is largely conserved between organoids and human tissue across developmental stages. These findings indicate that retinal organoids can serve as a viable model in which to dissect the function of specific noncoding regulatory elements. As a proof of concept, we perturbed a phylogenetically conserved enhancer at 5q14.3 that is accessible in human retinal progenitor cells and mature Müller glia, and that has been associated with retinal disorders such as Macular Telangiectasia Type 2 (MacTel) and Age-related Macular Degeneration (AMD) as well as related phenotypes including increased retinal vascular caliber and macular thickness (Bonelli et al., 2021; Gao et al., 2019; Han et al., 2020; Ikram et al., 2010; Scerri et al., 2017). To determine the role of this enhancer in gene regulation at this locus, we profiled cell-class-specific gene expression in enhancer knockout and isogenic control organoids over the course of retinal neurogenesis. These studies identify the long noncoding RNA *LINC00461*, the primary transcript of miR-9-2, as the principal target regulated by this enhancer. Deletion of this enhancer results in a decrease in LINC00461 expression, a de-repression of miR-9-2 targets, a concerted delay in the progression of retinal cell class specification, and a reduction in the relative number of rod photoreceptors, demonstrating a role for this enhancer in retinal development. Furthermore, regulation of miR-9-2 by the 5q14.3 enhancer plays a distinct role in more mature retinal organoids, where it controls expression of cell adhesion genes, metabolic function, and glial-vascular signaling molecules in Müller glia. These studies not only provide a rich resource for studying CREs over the course of human retinal development, but also demonstrate the utility of human-derived retinal organoids as a means for studying the function of disease-relevant noncoding regulatory elements.

## RESULTS

### Single-nucleus characterization of chromatin accessibility in the developing and mature human retina

To characterize chromatin accessibility at the single-cell level across human retinal development, we performed snATAC-seq on human retinal tissue at 6 developmental timepoints and in three adult donors. Adult human retinas were collected from two female donors (50 and 52 years old) and one male donor (54 years old). Developing retinal tissue was collected at 7, 8, 10, 11, 16, and 18 gestational weeks (gw) (Figure 1A-B). We generated and sequenced libraries for each sample, then integrated and analyzed the resulting datasets with the ArchR package (Granja et al., 2021). In parallel to these experiments, we performed snRNA-seq on the same pool of nuclei from each sample, analyzed the resulting data, and integrated the snRNA-seq and snATAC-seq datasets. The number of nuclei recovered and quality control data can be found in Table S1. We were thus able to use the gene expression data to aid in defining specific cell classes based on known retinal marker genes (Table S2). Within the resulting dataset, nuclei from 7 gw tissue are mostly clustered independently, and are predominantly comprised of early retinal progenitor cells (RPCs) and retinal ganglion cell (RGC) precursors (Figure 1B-C). Nuclei from 8, 10, and 11 gw tissue cluster together, and are comprised of late RPCs, developing RGCs, and a variety of retinal precursor cells. The abrupt transition in cell class development between 7 and 8 gw is supported by complementary studies describing a developmental epoch transition at this stage (Hoshino et al., 2017). Nuclei from 16 and 18 gw tissue cluster together, and still contain some late-stage RPCs, but are mostly made up of postmitotic, differentiating retinal cell classes. Finally, the adult tissues cluster together, and are comprised of the 7 major mature retinal cell classes: rod and cone photoreceptors; ganglion cells; horizontal cells; amacrine cells; bipolar cells; and Müller glia. To facilitate the use of these data in basic and disease research, we have assembled them, and additional data described below, into a searchable track hub via the University of California, Santa Cruz (UCSC) genome browser (https://tinyurl.com/CherryLab-SingleNuc-EyeBrowser).

**Figure 1:**
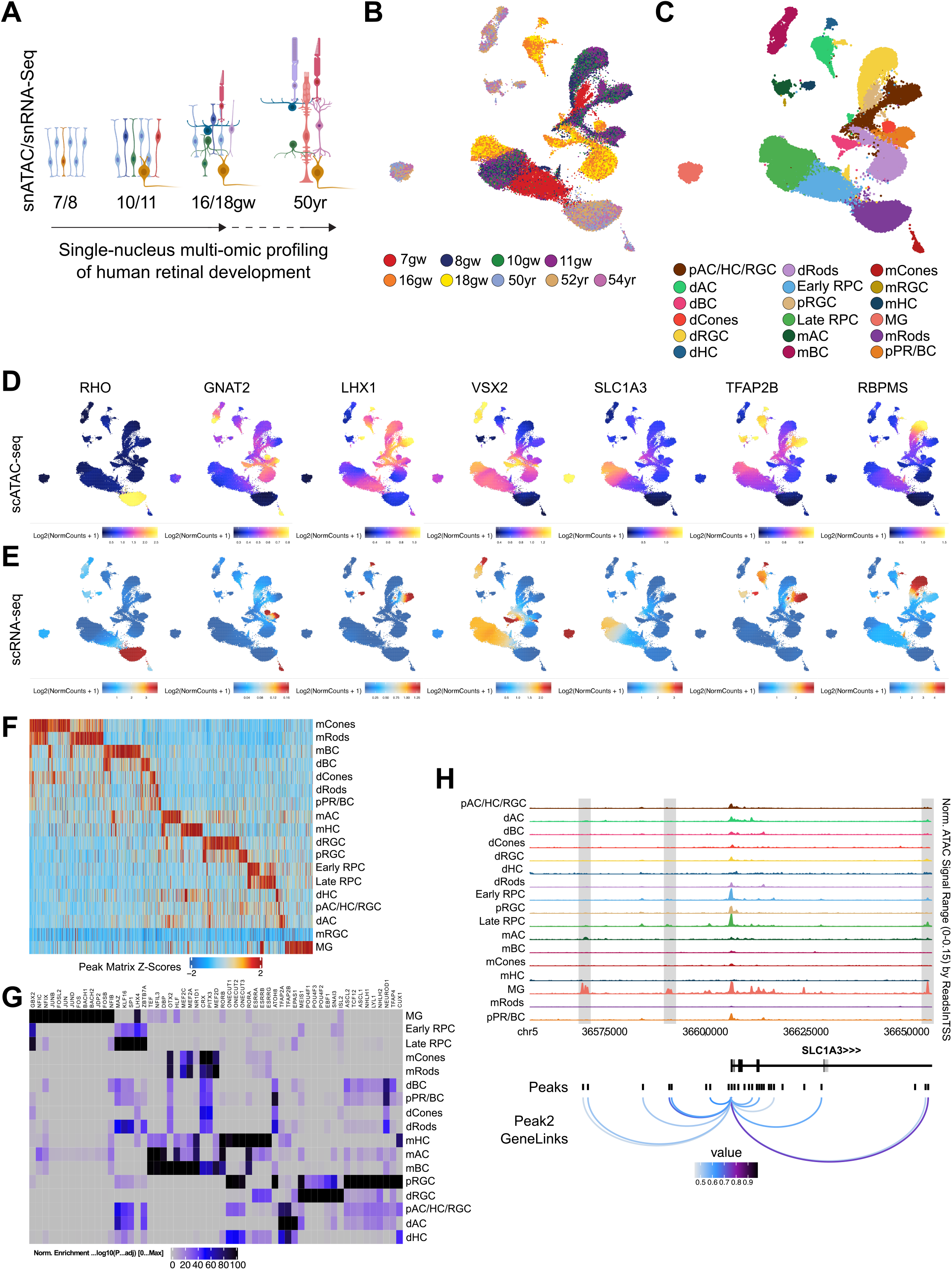
Single-cell chromatin accessibility profiling of developing and adult human retina. (**A**) Schematic of snATAC-seq/snRNA-seq experiments. Retinal tissue was harvested at the timepoints listed for the generation of single cell libraries. Gw = gestational week; yr = year. (**B**) UMAP of all cells from the snATAC-seq dataset colored by timepoint. (**C**) UMAP of all cells from the snATAC-seq dataset colored by cell class. Abbreviations: AC = amacrine cells; HC = horizontal cells; RGC = retinal ganglion cells; RPC = retinal progenitor cells; BC = bipolar cells; MG = Müller glia; PR = photoreceptors; p = precursor; d = developing; m = mature. (**D-E**) Feature plots showing the imputed gene score values (scATAC-seq, **D**) and the integrated gene expression values (scRNA-seq, **E**) for retinal cell class marker genes. (**F**) Heat map showing enrichment of DNA accessibility peaks in each cell class. (**G**) Heat map showing enrichment of transcription factor binding motifs in each cell class. (**H**) Peak-to-gene analysis at the *SLC1A3* locus. Top: Browser tracks indicating accessibility in each cell class around locus. Gray boxes highlight regions of high accessibility and linkage. Bottom: Called peaks and their associated linkage values with the *SLC1A3* promoter. Loops are colored by their linkage values. Darker colors indicate higher values. Scale (as shown) ranges from 0.5 to 1.

At the level of both chromatin accessibility and gene expression, these mature cell classes displayed the expected enrichment of cell-class-specific marker genes: *RHO* accessibility and expression are limited to mature rods; *GNAT2* is accessible and expressed in developing and mature cones; *LHX1* in developing and mature horizontal cells; *VSX2* in bipolar cells; *SLC1A3* in Müller glia; *TFAP2B* in amacrine and horizontal cells; and *RBPMS* in developing and mature RGCs (Figure 1D-E). We also performed trajectory analysis to examine changes in accessibility and expression over the course of the development and maturation of these distinct cell classes and observed the expected behavior of known early or late marker genes (Figure S1). *NRL*, which is required for rod development, increases drastically in both accessibility and expression over the course of pseudotime during rod development, whereas *NOTCH1*, which represses the rod fate, decreases in both accessibility and expression (Figure S1A). *ATOH7*, which is important for RGC survival during development (Brodie-Kommit et al., 2021), exhibits transient increases in accessibility and expression halfway through the course of RGC development (Figure S1G). Over the course of cone development, *NRL* expression remains low, as expected, but accessibility increases as cones mature, highlighting distinctions between accessibility and gene expression (Figure S1B).

These analyses allowed us to identify cell-class-enriched regions of accessibility, representing candidate CREs, for each of the major mature cell classes as well as those that are transiently enriched in discrete developmental cell states (Figure 1F). Within these regions, we further identified enriched families of transcription factor (TF) binding motifs in mature and developing cell classes. We filtered those motifs according to TFs expressed in those cell classes by snRNA-seq to identify the specific TFs that are most likely to target those motifs in each cell class (Figure 1G, Table S3). We observed enrichment of many expected motifs: the NFI family in late RPCs, Müller glia, amacrine and bipolar cells (Clark et al., 2019); OTX2/CRX in developing and mature photoreceptors and bipolar cells (Furukawa et al., 1997); the MEF2 family in mature retinal neurons (Andzelm et al., 2015); the POU4F factors in developing RGCs (Erkman et al., 1996; Gan et al., 1996); the ONECUT family in developing and mature horizontal cells and RGCs (Wu et al., 2012); and TFAP2 in amacrine and horizontal cells (Jin et al., 2015). Of note, developing and mature human cone peaks are enriched for the ISL2 motif, which has been shown to be more strongly expressed in human cones than in mouse cones (Lu et al., 2020). We also observed enrichment of unexpected motifs that have been less well described in specific retinal cell classes. The AP-1 factors (FOS/JUN) and the BACH family TFs were enriched in Müller glia and may reflect recent transduction of stimulus by these cells. The PAR bZIP family (TEF, DBP, HLF), which is known to accumulate in a circadian manner in several tissues (Gachon et al., 2004) but has not been extensively characterized in the retina, was enriched in Müller glia and in mature bipolar, amacrine, and horizontal cells.

Finally, we leveraged these snATAC-seq and snRNA-seq data to examine correlation between distal regions of accessibility that are likely enhancers and gene expression in all developing and mature cell classes (Table S4). As an example, at the *SLC1A3* locus, a gene important for Müller glia homeostasis, this analysis identified multiple regions of accessibility specific to Müller glia that were significantly associated with *SLC1A3* gene expression (Figure 1H; grey bars). This analysis is one important initial step toward solving the enduring problem of linking enhancers to their target genes that may be found at virtually any distance away from the enhancer.

### Conserved cis-regulatory landscapes between the human retina and retinal organoids

To evaluate hiPSC-derived retinal organoids as a system for studying cell-class-specific human retinal CREs in their native genomic environment, we performed snATAC-seq on retinal organoids over development (Figure 2A). We generated libraries for undifferentiated hiPSCs as well as for organoids that had been differentiating for 5-, 20-, and 28-weeks (2 biological replicates per timepoint). We chose these timepoints to approximate the stages that we profiled in the human retinal development experiment. We also performed snRNA-seq on nuclei from the same pools used for snATAC-seq and integrated these data via Seurat and ArchR (see Table S1 for quality control data). Within UMAP-space, the 5-week organoids cluster separately from their older counterparts and are comprised primarily of early RPCs, RGCs, and a variety of precursor cells. The 20- and 28-week organoids cluster together and contain all major mature retinal cell classes except for RGCs (Figure 2B-C), consistent with previous studies (Cowan et al., 2020; Sridhar et al., 2020). As in the human dataset, these mature retinal cell classes are enriched for their canonical marker genes at both the level of chromatin accessibility and gene expression (for instance, *RHO* expression in rods and *SLC1A3* in Müller glia; Figure S2A-B). These cell classes also display the expected patterns of accessibility and expression of known early and late marker genes over the course of development as assayed by trajectory analysis (Figure S2C-I).

**Figure 2:**
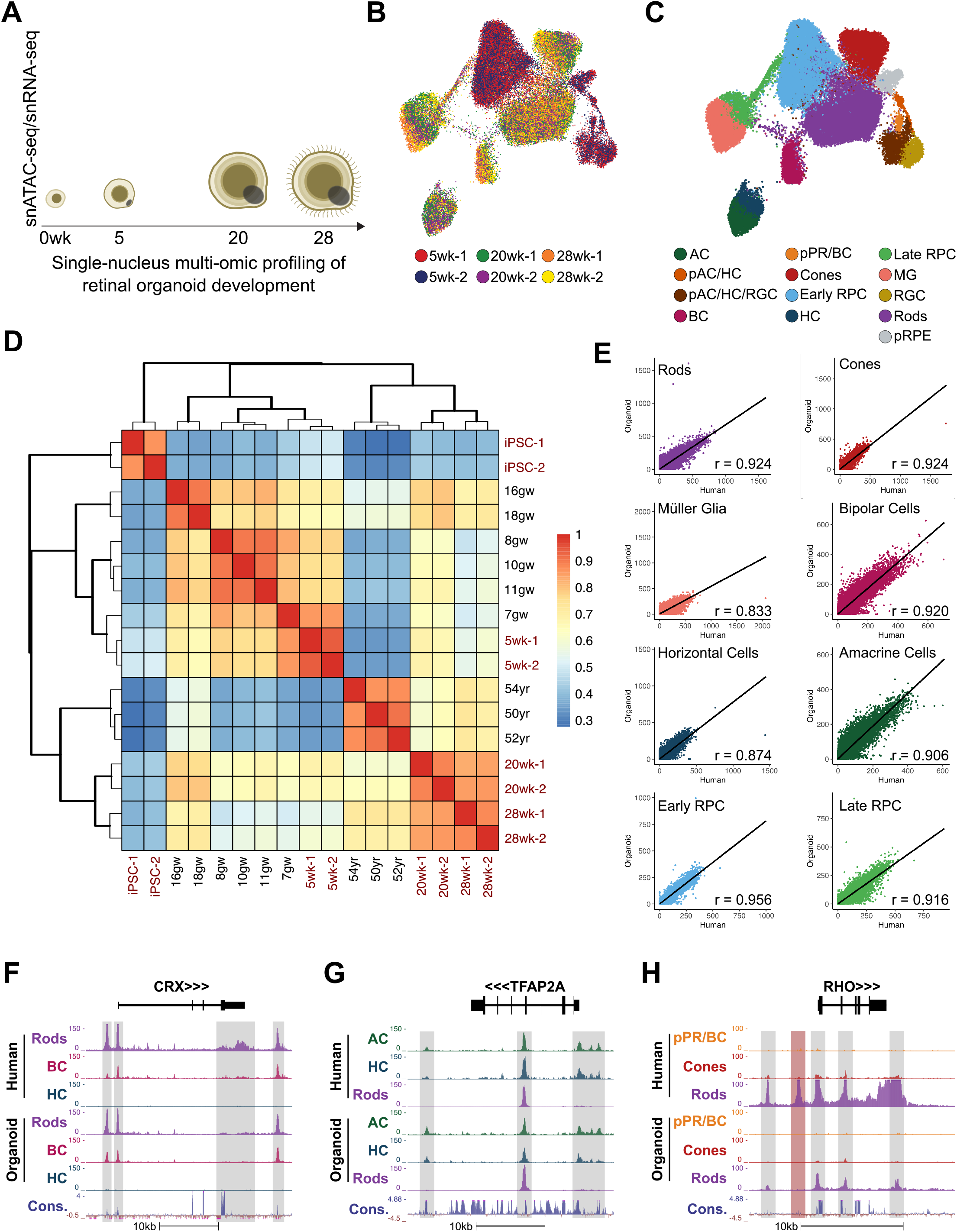
Chromatin accessibility landscape of human retina is conserved in human-derived retinal organoids. (**A**) Schematic of snATAC-seq/snRNA-seq experiments. Wk = weeks. (**B**) UMAP of all cells from the snATAC-seq dataset colored by timepoint. (**C**) UMAP of all cells from the snATAC-seq dataset colored by cell class. Abbreviations: AC = amacrine cells; PR = photoreceptors; BC = bipolar cells; RPC = retinal progenitor cells; HC = horizontal cells; MG = Müller glia; RGC = retinal ganglion cells; RPE = retinal pigment epithelium; p = precursor. (**D**) Heat map showing the correlation of pseudobulk DNA accessibility between human and organoid samples. Values are plotted as Pearson correlation coefficients. Red sample text indicates organoid tissue, and black text indicates human tissue. (**E**) Scatterplots depicting the correlation of peaks between organoid and human within individual retinal cell classes. Best-fit line is shown in black, along with the Pearson correlation coefficient. (**F-H**) Comparison of specific regions of accessibility between human and organoid cell classes at the *CRX* (**F**), *TFAP2A* (**G**), and *RHO* (**H**) loci. Gray boxes highlight accessible regions shared between tissue types. The Conservation track indicates conservation of regions across vertebrate species. The red box in (**H**) highlights a region of accessibility that is not shared between human and organoid rods.

To directly compare the chromatin accessibility landscape of the human retina and retinal organoids, we first correlated the patterns of DNA accessibility between each sample using a pseudobulk strategy (Figure 2D). Undifferentiated hiPSCs correlate strongly with themselves but poorly with any other timepoint. 5-week organoids have the highest correlation with 7 gw human tissue. 20-week organoids correlate with developing retinas, most strongly at 16-18 gw. 28-week organoids correlate more strongly with late gestational (16-18 gw) and adult retinas than with earlier timepoints. We next performed a pairwise correlation between human and organoid samples in a cell-class-specific manner, comparing all the major mature retinal cell classes (with the exception of RGCs due to low numbers) as well as early and late RPCs. All of these cell classes display a high degree of correlation between the retina and organoids, with a Pearson correlation coefficient ranging from 0.833 between Müller glia to 0.956 between early RPCs (Figure 2E).

Similarities in the epigenomic landscapes of human retinas and organoids can be seen at the level of individual loci as well. For instance, there are numerous regions of accessibility surrounding the *CRX* locus that are present in rods and bipolar cells, but not in horizontal cells (Figure 2F, gray rectangles). This pattern is conserved between the human and organoid cell classes. At the *TFAP2A* locus, there are regions upstream and downstream of the locus that are accessible in horizontal and amacrine cells, but not in rods, and another site that is accessible in all three cell classes (Figure 2G, gray rectangles). This pattern of accessibility is also conserved in organoids. However, we do observe differences in accessibility patterns as well. While accessibility around the *RHO* locus is largely conserved between tissues (Figure 2H, gray rectangles), there is a site upstream of the *RHO* promoter that is accessible in mature human rods but not in organoid rods (Figure 2H, red rectangle). This region has previously been characterized as a likely *RHO* enhancer (Cherry et al., 2020), and its absence in organoid rods could be a determining factor in the more modest levels of RHO expression in organoids compared to adult human retinas. CREs like this may represent opportunities to further optimize organoid models by targeted activation of these enhancers. Despite these differences, the high degree of global correlation of chromatin accessibility at the cell class level between tissues indicates that human retinal organoids are a reasonably good model for studying the function of retinal CREs. This comparison provides a valuable resource for any future studies investigating human retinal CREs.

### Disease-associated loci and DNA accessibility in the human retina and model systems

Genome-wide association studies (GWAS) frequently identify disease association with non-coding regions of the genome, suggesting that genetic variants within CREs might contribute to disease phenotypes (Maurano et al., 2012). To gain insight into the potential contribution of common variants within CREs to retinal disease, we examined the intersection between disease-associated loci from two retinal disease GWAS studies and cell-class-specific regions of DNA accessibility in the human retina (Fritsche et al., 2016; Scerri et al., 2017) (Figure 3). In the first comparison, we focused on an intergenic region at 5q14.3 that has been associated with macular thickness, retinal vascular caliber, and AMD, and is one of the most highly associated peaks with MacTel (Bonelli et al., 2021; Gao et al., 2019; Han et al., 2020; Ikram et al., 2010; Scerri et al., 2017). A number of DNA accessibility peaks fall within this locus (Figure 3A). Of note, the lead sentinel single nucleotide polymorphism (SNP) (rs17421627) for this associated region intersects an enhancer that we and others have previously shown 1) to be sufficient to drive gene expression in the vertebrate retina, and 2) that the risk SNP significantly decreases the activity of this enhancer (Cherry et al., 2020; Madelaine et al., 2018) (Figure 3A, gray rectangle). In this current analysis, we found that the cell-class-specific pattern of DNA accessibility at this enhancer is well conserved between the human retina and organoids (Figure 3C, gray rectangle). In both retina and organoids this enhancer is predominantly accessible in early and late RPCs and in Müller glia. This cell-class-specificity is consistent with the phenotype of MacTel which, is characterized by the progressive loss of Müller glia (Powner et al., 2013).

**Figure 3:**
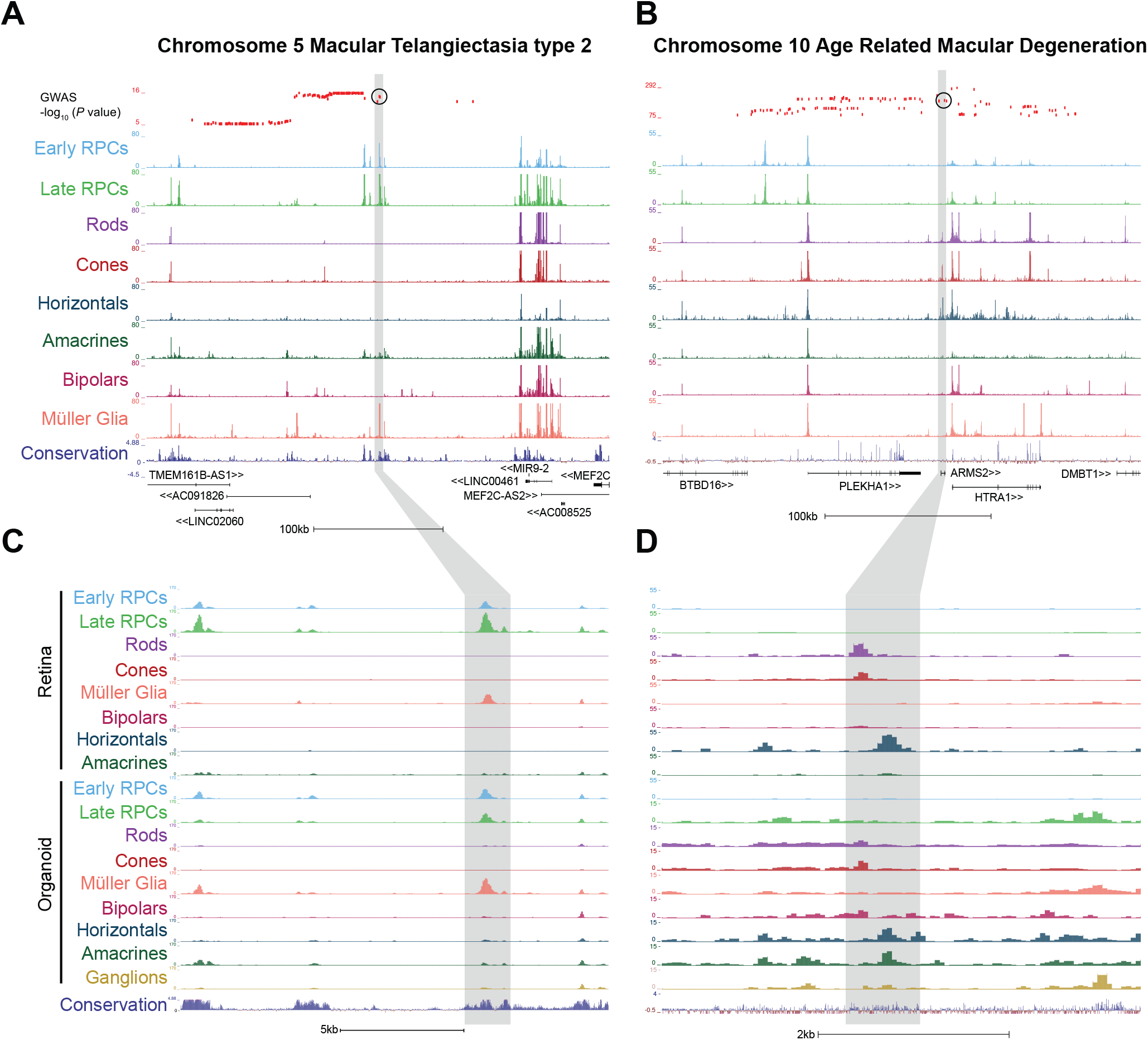
Overlap of GWAS loci and chromatin accessibility in human and organoid tissue. (**A-B**) Alignment of GWAS hits with chromatin accessibility in individual human retinal cell classes at the 5q14.3 (**A**) and 10q26 (**B**) loci. Red ticks indicate the strength of the SNP association. Sentinel SNPs are circled in black. Gray bars highlight the regions showcased in C-D. The Conservation tracks indicates conservation of regions across vertebrate species (**C-D**) Comparison of chromatin accessibility between human and organoid cell classes around the regions highlighted in A-B. The Conservation tracks indicates conservation of regions across vertebrate species

In the second comparison, we focused on a region at 10q26 that is one of the strongest loci linked to AMD risk (Fritsche et al., 2016). We found that this locus also contains a number of putative CREs and that the sentinel SNP (rs3750846) is proximal to a region of accessibility within the intron of the *ARMS2* locus (Figure 3B, gray rectangle). The sequence inclusive of this accessibility peak has previously been proposed to act as a distal regulatory element, and risk SNPs within this area may contribute to misregulation of nearby target genes and therefore AMD pathobiology (Liao et al., 2017). We find that this area actually contains two distinct peaks. One peak is shared between rod and cone photoreceptor cells and the other is enriched in horizontal cells of the human retina (Figure 3D). These patterns of accessibility can be observed in the same cell classes in human retinal organoids. Although it is currently unknown what the significance of these regions is to MacTel and AMD, the demonstration that the lead sentinel SNPs at both the 5q14.3 and the 10q26 GWAS loci intersect cell-class-specific accessibility peaks helps to prioritize which cell classes may be primarily affected in these diseases and which non-coding regulatory elements to prioritize for functional investigations.

Taking a complementary approach to this GWAS analysis, we also sought to determine the ability of organoids to model the CREs of known retinal disease genes. We reasoned that CREs of known retinal disease genes may play important roles in development and maintenance of retinal cell types and may harbor non-coding variants that contribute to disease. There are over 260 genes to date that have been implicated in human visual disorders (RetNet; https://sph.uth.edu/retnet/). For each of these genes we first identified candidate regulatory elements based on the correlation between single-cell gene expression with single-cell DNA accessibility patterns in the developing and adult human retina (Table S4, S5). The vast majority of these CREs are enriched in rod and cone photoreceptor cells, with a smaller number enriched within other cells classes such as Müller glia, amacrine cells, and RPCs (Figure S3A). Next, we compared DNA accessibility at these candidate CREs in the human retina to regions of accessibility that are associated with expression of the same genes in organoids. Using unsupervised clustering, we observed that the cell-class-specific patterns of DNA accessibility at candidate disease-gene-associated CREs is well conserved between human retinas and organoids. These observations further underscore the value of human retinal organoids as a system to model the function of disease-gene-associated CREs.

Although we present evidence justifying human retinal organoids as a powerful model for studying the function of disease-associated CREs, certain retinal diseases affect aspects of retinal biology which are currently difficult to model *in vitro* in retinal organoids. We therefore sought to benchmark these human disease-gene-associated CREs with the most commonly used *in vivo* model, the developing and adult mouse retina (Figure S3B). We utilized snATAC-seq data generated by Lyu, Blackshaw, and colleagues (Lyu, 2021) and performed a similar comparison between human retinal CREs and mouse retinal CREs. We found that unsupervised clustering of snATAC-signal at disease-gene-associated CREs also paired homologous mouse and human cell classes together, suggesting that mice may serve as a complementary system to model these CREs in an *in vivo* context despite the great evolutionary distance between humans and mice. Indeed, the fact that the likely function of these CREs has been preserved over evolutionary time underscores the importance of individual CREs in visual species.

### A disease-associated enhancer at 5q14.3 regulates the primary transcript of miR-9-2

As a proof of concept that these single nucleus multi-ome studies can help direct investigations into the function of human retinal CREs in development and disease we focused our efforts on the 5q14.3 enhancer that we described above (which we refer to as e5q14.3). To determine the function of this enhancer in a human retina-like system, we excised the highlighted region in Figure 3C in hiPSCs via CRISPR editing and differentiated these cells into human enhancer knockout (KO) retinal organoids. We then performed snRNA-seq on knockout organoids and their isogenic controls at 5, 12, 20, and 28 weeks of development and examined changes in single-nucleus gene expression between the two genotypes (Figure 4A). For each timepoint we assayed multiple biological replicates from independent clones for each genotype (5wk, n=2; 12, 20, and 28wks, n=3). To account for variability between individual organoids, each biological replicate represents a pool of 4-6 organoids. Following analysis and visualization with Seurat, we observed all of the expected major retinal cell classes in both enhancer knockout and control organoids (Figure 4B).

**Figure 4:**
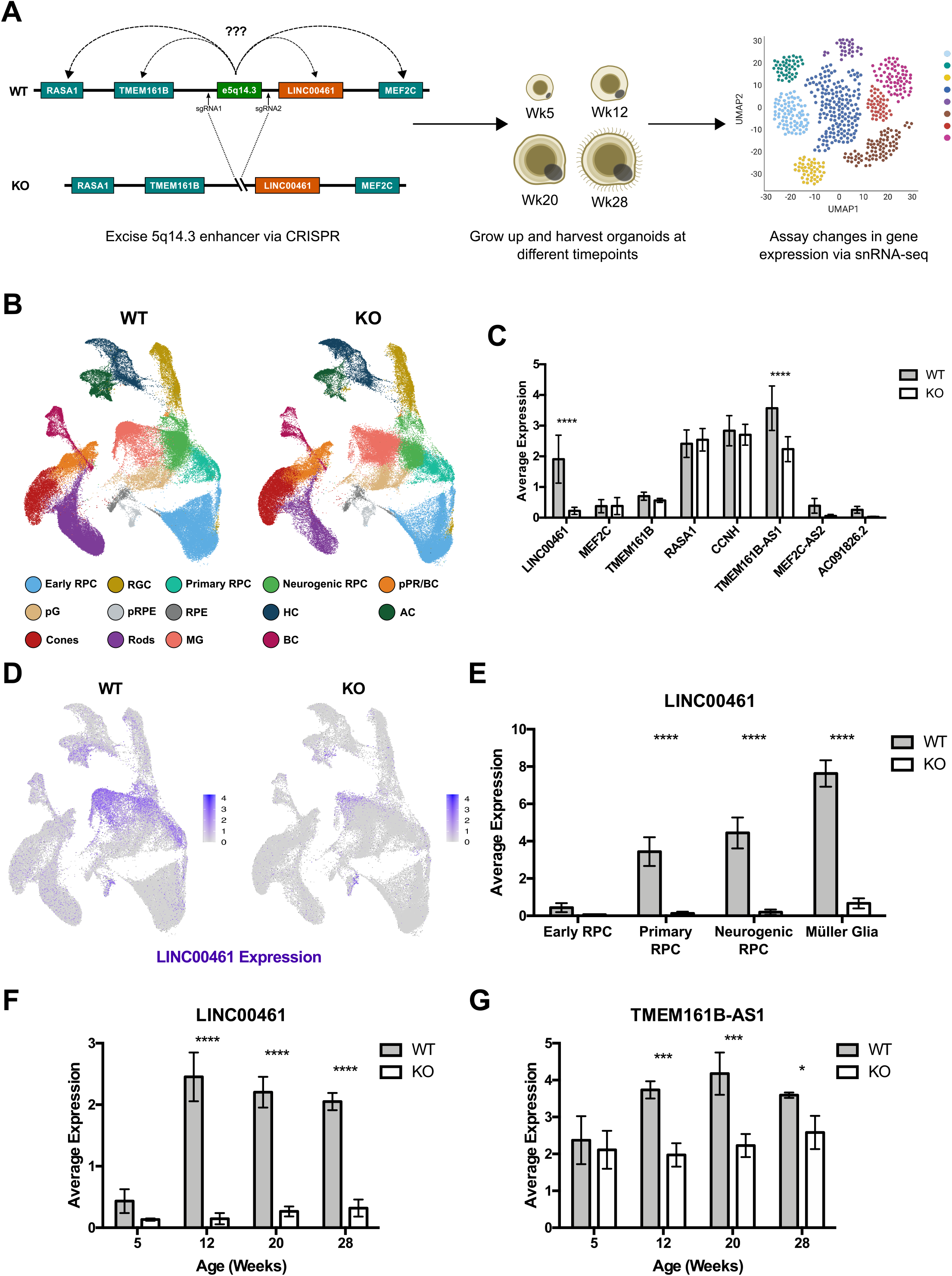
The 5q14.3 enhancer regulates *LINC00461* in retinal organoids. (**A**) Schematic of e5q14.3-deletion experiment. CRE was excised in iPSCs, which were then differentiated into retinal organoids and harvested at the timepoints indicated for the generation of snRNA-seq libraries. (**B**) UMAPs of all cells from the snRNA-seq dataset split by wildtype (WT) and enhancer knockout (KO) and colored by cell class. Abbreviations: RPC = retinal progenitor cells; RGC = retinal ganglion cells; PR = photoreceptors; BC = bipolar cells; G = glia; RPE = retinal pigment epithelium; HC = horizontal cells; AC = amacrine cells; MG = Müller glia; p = precursor. (**C**) Average expression values of the genes within and adjacent to the e5q14.3 TAD in WT (gray bars) and KO organoids (white bars). n = 11 samples. Values are reported as mean ± SD. Two-way ANOVA, Sidak post-test. **** = p ≤ 0.0001. (**D**) Feature plots showing expression of *LINC00461* in WT and KO organoids. Darker colors indicate higher expression. (**E**) Average expression values of *LINC00461* between WT (gray) and KO (white) organoids specifically within RPCs and Müller glia. Early RPC: n = 2 samples. Primary and Neurogenic RPCs, Müller glia: n = 3 samples. Values are reported as mean ± SD. Two-way ANOVA, Sidak post-test. **** = p ≤ 0.0001. (**F-G**) Average expression values of *LINC00461* (**F**) and *TMEM161B-AS1* (**G**) between WT (gray) and KO (white) organoids at each timepoint. 5 weeks: n = 2 samples. 12, 20, and 28 weeks: n = 3 samples. Values are reported as mean ± SD. Two-way ANOVA, Sidak post-test. * = ≤ 0.05; *** = p ≤ 0.001; **** = p ≤ 0.0001.

To begin to understand the function of this enhancer, we first sought to determine its direct target gene(s). Because CREs can regulate genes at variable distances, we limited our genomic inquiry to the presumptive topologically associating domain (TAD) containing e5q14.3 (which we provisionally determined to be chr5:87,781,318-88,691,140 (hg38) based on Hi-C and CTCF-binding data; Figure S4) (Cherry et al., 2020; de Bruijn et al., 2020) and the TAD-adjacent genes *MEF2C*, *RASA1*, and *CCNH* that have been suggested to play a role in 5q14.3-associated phenotypes. Of all the TAD genes that were present in our dataset, only the long noncoding RNA *LINC00461* and the anti-sense transcript *TMEM161B-AS1* were significantly downregulated within the global knockout dataset (Figure 4C-D). Decreased expression of both *LINC00461* and *TMEM161B-AS1* began at 12 weeks and persisted throughout the later timepoints (Figure 4F, G respectively). The proportionate decrease in *LINC00461* expression was far greater than that of *TMEM161B-AS1*. While *TMEM161B-AS1* has no known biological function, *LINC00461* is understood to be the primary transcript for the conserved, developmentally important microRNA miR-9-2 (Oliver et al., 2015). Thus, these results suggest that *LINC00461* is the principal functional target of e5q14.3, although it can regulate other transcripts as well. This finding, together with the recent work of Madelaine and colleagues in zebrafish, suggests that regulation of the miR-9-2 primary transcript by this conserved enhancer may be an ancient mechanism in vertebrates (Madelaine et al, 2018). Lastly, given that activity of e5q14.3 was enriched in RPCs and Müller glia, we looked to determine if e5q14.3 is required for LINC00461 expression in these specific cell classes. Indeed, our snRNA-seq data demonstrated significant decreases in LINC00461 expression in primary and neurogenic RPCs and in Müller glia, indicating that these cell classes may use e5q14.3 regulation of LINC00461 for specific biological functions (Fig. 4E). In contrast, there was no significant change in LINC00461 expression in early RPCs. We therefore focused on these progenitor populations and glia for downstream analyses.

### The 5q14.3 enhancer regulates the timing of retinal cell class specification

Given that e5q14.3 regulates expression of LINC00461 in primary and neurogenic RPCs, we sought to determine what role this enhancer might have in retinal organoid development. Notably, we observed significant differences in the proportions of distinct cell classes during the development of enhancer KO and control organoids (Figure 5A-D). The most significant difference was the decrease in the proportion of rod photoreceptors in enhancer KO organoids. Immunostaining of photoreceptors in KO and control organoids showed that there is a noticeable thinning of the putative outer nuclear layer (ONL), supporting a decrease in rods, in enhancer KO organoids (Figure 5B-C). Quantification of other retinal cell classes from early to late organoid development showed other significant differences. At 12 weeks, an early stage of development, there were significantly more neurogenic and primary RPCs in KO organoids. Furthermore, while photoreceptor/bipolar cell precursors and horizontal cells were initially depleted in KO organoids, their proportions were later over-represented by 20 weeks. Finally, at late stages of development (20-28 weeks), cones and Müller glia are over-represented, while rods and bipolar cells are depleted in KO organoids. By 28 weeks however, bipolar cells catch up in the KO versus control organoids. Altogether, these shifts in cell-class proportions suggest that the timing of retinal neuronal cell class specification is delayed in KO organoids (Figure 5E). Additionally, the timing of Müller glial cell specification appeared normal in enhancer KO organoids, suggesting an underlying uncoupling of the timing of neurogenesis and gliogenesis.

**Figure 5:**
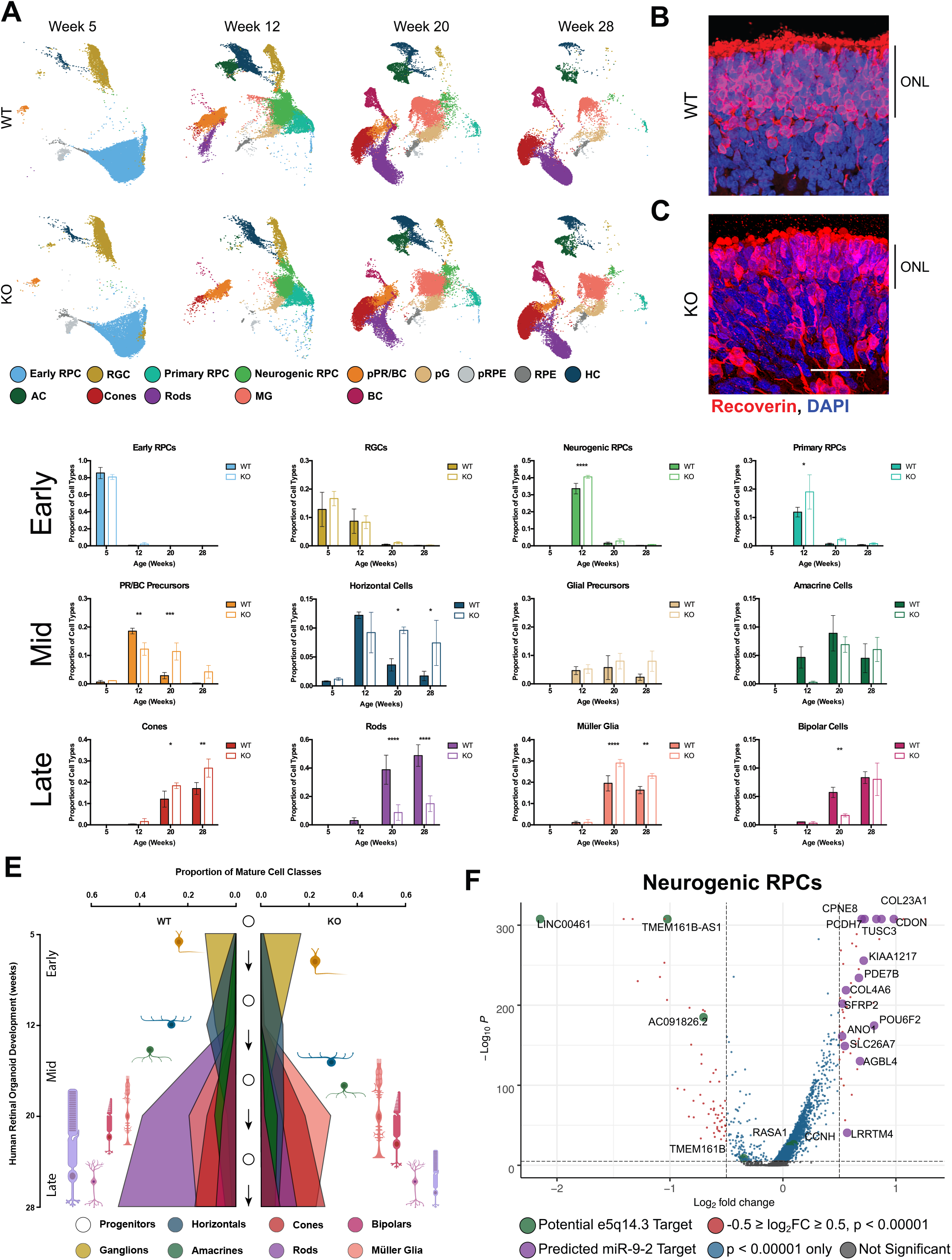
The 5q14.3 enhancer regulates timing of neurogenesis in retinal organoids. (**A**) UMAPs of all cells from the snRNA-seq dataset split by genotype and by timepoint and colored by cell class. Abbreviations: RPC = retinal progenitor cells; RGC = retinal ganglion cells; PR = photoreceptors; BC = bipolar cells; G = glia; RPE = retinal pigment epithelium; HC = horizontal cells; AC = amacrine cells; MG = Müller glia; p = precursor. (**B-C**) Maximum projections of sections from WT (**B**) and KO (**C**) organoids immunostained with an anti-Recoverin antibody at 20 weeks. Recoverin-positive cells are shown in red, and DAPI-labeled nuclei are shown in blue. The size of the outer nuclear layer (ONL) is labeled on the righthand side of each image. Scale bar in (**C**) = 50 µm. (**D**) Proportions of all cell classes in WT (filled bars) and KO (unfilled bars) organoids at each timepoint. 5 weeks: n = 2 samples. 12, 20, and 28 weeks: n = 3 samples. Values are reported as mean ± SD. Two-way ANOVA, Sidak post-test. * = ≤ 0.05; ** = ≤ 0.01; *** = p ≤ 0.001; **** = p ≤ 0.0001. (**E**) Schematic showing the proportion of cell classes (x-axis) in WT and KO organoids over developmental time (y-axis). (**F**) Volcano plot depicting all differentially expressed genes in KO neurogenic RPCs. Dashed lines indicate significance thresholds: a significant p-value was set at p < 0.00001 (or -log(p) of 5) and significant log2 fold changes were set at ± 0.5. Gray dots indicate genes with no significant change. Blue dots indicate genes with a significant p-value but not a significant fold change. Red dots indicate genes with both a significant p-value and significant fold change. Green dots highlight genes that are potential e5q14.3 targets. Purple dots highlight genes that are predicted miR-9-2 targets. Green and purple dots are labeled with their gene names.

One explanation for these observed differences in developmental timing could be a change in retinal progenitor cell state driven by altered gene expression. We therefore compared gene expression in primary and neurogenic RPCs in enhancer KO and control organoids (Figures 5F, S5A, Table S6). In both classes, LINC00461 is the most significantly downregulated gene. Other potential targets of e5q14.3 are less downregulated or do not reach significance. Of all of the significantly dysregulated genes, we observed that more genes were upregulated in KO progenitors, consistent with the effects of the loss of a miRNA. To determine if miR-9 targets are de-repressed in primary or neurogenic RPCs, we used TargetScan (Agarwal et al., 2015) to identify candidate genes that are predicted to be repressed by miR-9 (including both miR-9-3 and miR-9-5 predicted targets). Out of the upregulated genes in progenitors, many are predicted to be targets of miR-9-2 including *TUSC3*, *COL23A1*, *CPNE8*, *CDON*, *PCDH7*, and others (Figures 5F, S5A, purple dots). However, the change in developmental timing is likely not caused by any single gene, but rather is the net consequence of misregulation of a concerted gene regulatory network mediated by miR-9-2. We therefore performed gene set enrichment analysis (GSEA) of differentially expressed genes in knockout RPC populations which revealed an enrichment of upregulated genes in pathways involved in cell division (Myc and E2F signaling, G2M checkpoint) and in metabolic pathways that facilitate the high energy demand of cell division (mTORC1 signaling, oxidative phosphorylation) (Figures S5B-C, S7), suggesting that cell cycle regulation may be altered in knockout RPCs. The Notch signaling pathway, which regulates cell fate during retinal neurogenesis (Jadhav et al., 2006a; Jadhav et al., 2006b), is also associated with upregulated genes (Figures S5B-C, S7). These data suggest that e5q14.3 regulates miR-9-2 expression in primary and neurogenic RPC populations, which may influence the timing of retinal development leading to a change in the distribution of mature retinal cell types.

The differences we observe may also be influenced by a delay in progenitor cells giving rise to postmitotic daughter cells. To address this possibility, we performed birthdating experiments using an EdU pulse-chase approach. Since the rod population was most impacted, we birthdated cells at a timepoint when rods are arising in control organoids. First, we administered a 24-hour pulse of EdU to control and KO organoids at week 16 and either harvested them immediately afterwards or allowed them to grow to week 22, when cells should be post-mitotic and express mature retinal cell markers (Figure 6A). At week 16 we observed significantly more EdU-labeled cells in KO organoids (Figure 6B-B’), indicating an increase in the number of progenitors, consistent with cell distributions seen at week 12. We next examined the identity of week 16 labeled EdU-positive cells after we let them develop to week 22 by co-staining with markers for photoreceptors and Müller glia, the predominant cell classes born at week 16. Cells that were born within one or two cell cycles of the initial pulse at week 16 should be identifiable as heavily EdU-labeled differentiated cells by week 22. We observed a slight but nonsignificant decrease in the proportion of EdU-positive photoreceptors in KO organoids and observed no difference in the Müller glia (Figure 6C-C’ and Figure 6D-D’, respectively). However, by week 22 the KO organoids contained significantly fewer total EdU-positive cells (Figure 6E-E’), indicating that either fewer birthdated cells were surviving in KO organoids or fewer progenitors exited the cell cycle and therefore the EdU label was diluted out over subsequent cell divisions. To assay cell death, we performed TUNEL staining at weeks 16 and 20 and found no evidence of increased cell death in the KO organoids (Figure S6A-B’). We next administered a 24-hour EdU pulse at week 20, when there is less proliferation in control organoids, and harvested organoids immediately afterward (Figure 6A). At this time point we observed that KO organoids still had significantly greater numbers of proliferating cells (Figure 6F-F’) suggesting that, in the absence of e5q14.3, RPCs remain in the cell cycle for a longer window of organoid development, and thus dilute out their EdU signal between weeks 16 and 22. Altogether, these data demonstrate that the loss of e5q14.3 alters the proliferative capacity of RPCs, resulting in a delay in the production of later-born retinal cell classes.

**Figure 6:**
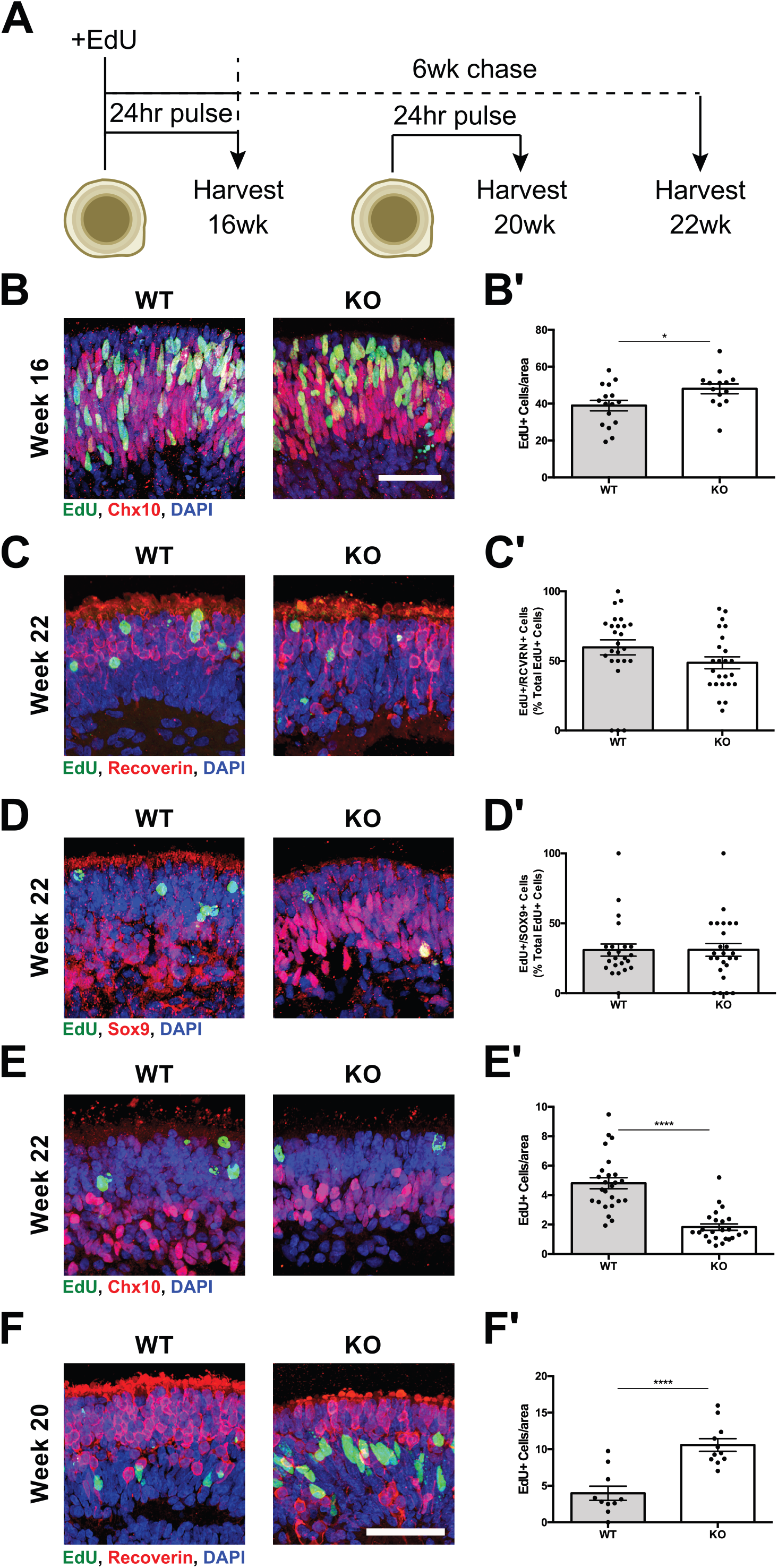
Deletion of the 5q14.3 enhancer in organoids results in delayed cell cycle exit of retinal progenitor cells. (**A**) Schematic of EdU pulse-fix/pulse-chase experiments. EdU was administered to WT and KO organoids for 24 hours at week 16, and organoids were either harvested immediately after the pulse or at week 22. A separate batch of organoids was administered EdU for 24 hours at week 20 and harvested immediately afterward. (**B**) Maximum projections of sections from WT and KO organoids following EdU treatment at week 16. EdU-positive cells are shown in green, Chx10-positive cells are shown in red, and DAPI-labeled nuclei are shown in blue. Scale bar = 50 µm. (**B’**) Quantification of EdU-positive cells in WT (gray) and KO (white) organoids at week 16. n = 16 organoids (WT); n = 14 organoids (KO). Values are reported as mean ± SEM. Student’s unpaired t test. * = p ≤ 0.05. (**C**) Maximum projections of sections from WT and KO organoids harvested at week 22 and immunostained for photoreceptors. EdU-positive cells are shown in green, Recoverin-positive cells are shown in red, and DAPI-labeled nuclei are shown in blue. (**C’**) Quantification of the percentage of EdU-positive photoreceptors in WT (gray) and KO (white) organoids at week 22. n = 25 organoids (WT); n = 24 organoids (KO). Values are reported as mean ± SEM. Student’s unpaired t test. p = 0.1160. (**D**) Maximum projections of sections from WT and KO organoids harvested at week 22 and immunostained for Müller glia. EdU-positive cells are shown in green, Sox9-positive cells are shown in red, and DAPI-labeled nuclei are shown in blue. (**D’**) Quantification of the percentage of EdU-positive Müller glia in WT (gray) and KO (white) organoids at week 22. n = 23 organoids (WT); n = 25 organoids (KO). Values are reported as mean ± SEM. Student’s unpaired t test. p = 0. 9835. (**E**) Maximum projections of sections from WT and KO organoids following EdU treatment at week 16 and harvested at week 22. EdU-positive cells are shown in green, Chx10-positive cells are shown in red, and DAPI-labeled nuclei are shown in blue. (**E’**) Quantification of EdU-positive cells in WT (gray) and KO (white) organoids harvested at week 22. n = 25 organoids (WT); n = 24 organoids (KO). Values are reported as mean ± SEM. Student’s unpaired t test. **** = p ≤ 0.0001. (**F**) Maximum projections of sections from WT and KO organoids following EdU treatment at week 20. EdU-positive cells are shown in green, Recoverin-positive cells are shown in red, and DAPI-labeled nuclei are shown in blue. Scale bar = 50 µm (also applies to **C-E**). (**F’**) Quantification of EdU-positive cells in WT (gray) and KO (white) organoids at week 20. n = 10 organoids (WT). n = 11 organoids (KO). Values are reported as mean ± SEM. Student’s unpaired t test. **** = p ≤ 0.0001.

### The 5q14.3 enhancer regulates a distinct network of gene expression in Müller glia

Given that e5q14.3 also regulates expression of LINC00461 in post-mitotic Müller glial cells (Figure 4E), and that many phenotypes associated with 5q14.3 variants could potentially have an origin in Müller cell dysfunction, we lastly sought to determine what role this enhancer might have in maintaining the gene regulatory networks of Müller glia. Immunostaining of Müller glia shows the presence and morphology of Müller glia in control and enhancer KO organoids (Figure 7A). We identified differentially expressed genes in KO Müller glia compared to isogenic controls (Figure 7B, Table S6). Similar to the late RPCs, *LINC00461* is the most significantly downregulated gene. KO Müller glia also exhibit significant decreases in other TAD genes, such as *TMEM161B-AS1*, *AC09186.2*, and *MEF2C-AS2* (Figure 7B, green dots). We observed increased expression of a number of predicted miR-9-2 target genes in knockout Müller glia (Figure 7B, purple dots), suggesting that miR-9-2 function is also impaired in Müller glia. Some of these de-repressed targets are shared with KO RPCs while others are distinct, such as cell adhesion molecules *CNTN3*, *4*, and *5*, the transcription factor *MEIS1*, and the AMD-associated gene *PLEKHA1*. Interestingly, we also observe decreased expression of genes related to vascular function, such as *FLT1*, *SEMA3A*, and *COL18A1* (Figure 7B, yellow dots). VEGFA is also downregulated in enhancer KO Müller glia (Figure S7), consistent with previous studies of the miR-9 family (Madelaine et al., 2017), albeit to a lesser degree compared to these other vascular-related molecules. Notably, *COL18A1* is the causal gene of Knobloch syndrome which is associated with several ocular abnormalities. Müller glia are intimately associated with retinal vasculature (Vecino et al., 2016), so these results suggest that the loss of e5q14.3 may lead to the vascular phenotypes associated with the 5q14.3 locus through dysregulated glial-vascular signaling. Globally, GSEA of differentially expressed genes in KO Müller glia indicate that upregulated genes are associated with many of the same pathways enriched in RPCs, such as cell division pathways (Myc and E2F signaling, G2M checkpoint) and Notch signaling (Figure 7C). However, in contrast to the RPCs, many metabolic pathways (mTORC1 signaling, oxidative phosphorylation, adipogenesis) are highly associated with downregulated genes. Most remarkably, genes associated with the hypoxia pathway also show this opposing behavior in Müller glia compared to RPCs (Figure S7). Overall, these data suggest a role for e5q14.3 in regulating miR-9-2 and its direct and indirect targets in Müller glia to support Müller glia homeostasis. These observations are supported by an essential role for microRNAs, and the importance of the miR-9 family in particular, in the Müller glia of the mature mouse retina (Wohl et al., 2017).

**Figure 7:**
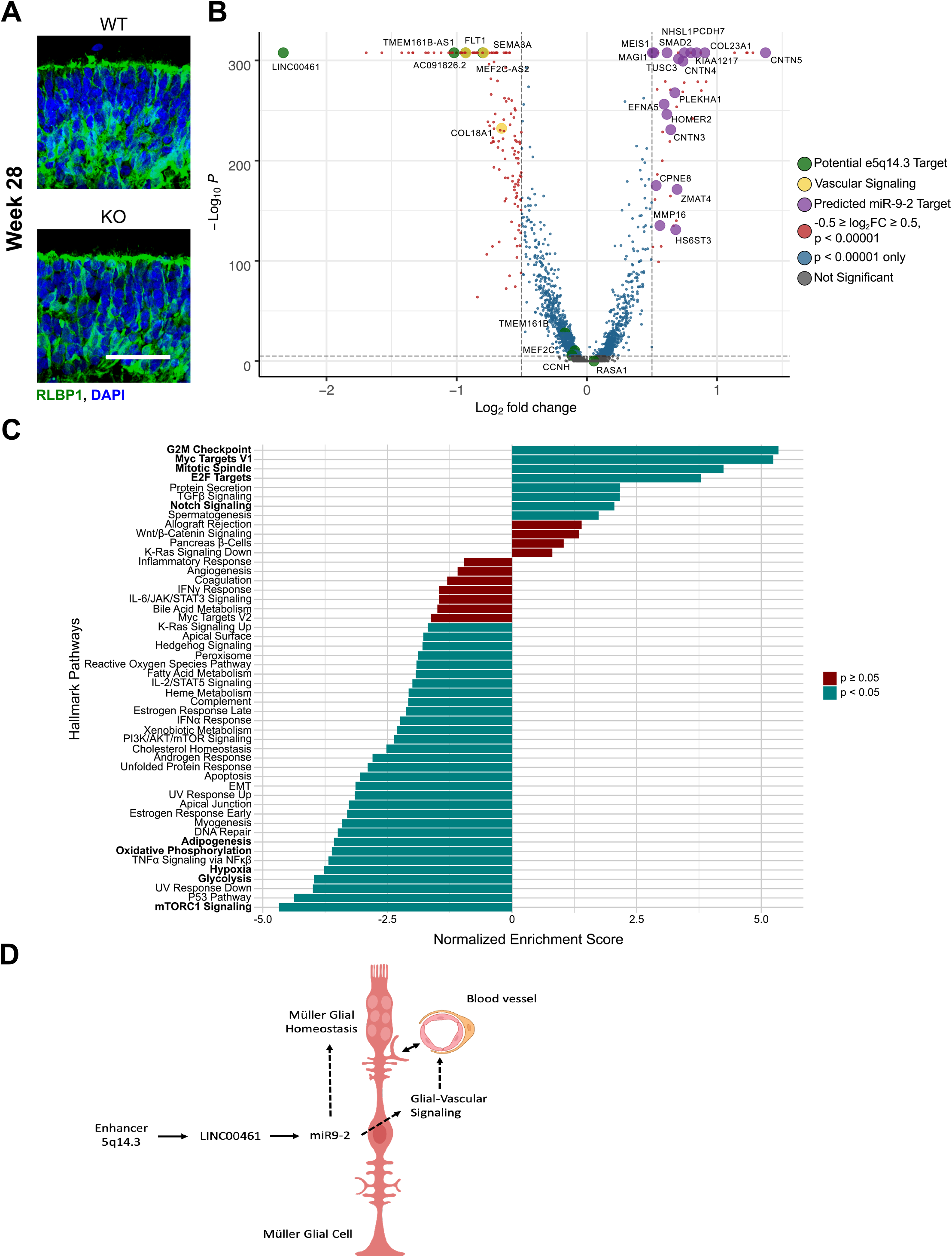
The impact of 5q14.3 enhancer deletion in Müller glia. (**A**) Maximum projections of sections from wildtype (WT) and knockout (KO) organoids immunostained with an anti-RLBP1 antibody at week 20. RLBP1-positive cells are shown in green, and DAPI-labeled nuclei are shown in blue. Scale bar = 50 µm. (**B**) Volcano plot depicting all differentially expressed genes in KO Müller glia. Dashed lines indicate significance thresholds: a significant p-value was set at p < 0.00001 (or -log(p) of 5) and significant log2 fold changes were set at ± 0.5. Gray dots indicate genes with no significant change. Blue dots indicate genes with a significant p-value but not a significant fold change. Red dots indicate genes with both a significant p-value and significant fold change. Green dots highlight genes that are potential e5q14.3 targets. Purple dots highlight genes that are predicted miR-9-2 targets. Yellow dots highlight genes that are involved in vascular signaling. Green, purple, and yellow dots are labeled with their gene names. (**C**) Gene set enrichment analysis of differentially-expressed genes in KO Müller glia. Normalized enrichment scores (NES) are plotted for all hallmark pathways. NES with an adjusted p-value of < 0.05 are shown in teal; NES with an adjusted p-value of ≥ 0.05 are shown in red. Pathways of interest are bolded. (**D**) Schematic of hypothesized function of e5q14.3 function in Müller glia. The enhancer regulates *LINC00461* expression and thus miR-9-2 expression, which in turn may regulate Müller glia homeostasis and/or signaling between Müller glia and the surrounding retinal vasculature.

## DISCUSSION

Cis-regulatory elements control essential gene expression needed for the development and maintenance of the human retina and have been found to be disrupted in distinct retinal disorders (Bhatia et al., 2013; Ghiasvand et al., 2011; Nathans et al., 1989). However, despite considerable recent progress, our current knowledge of human retinal CREs is incomplete and lacks cell-class-specific resolution (Aldiri et al., 2017; Cherry et al., 2020; Xie et al., 2020). This gap in knowledge prevents systematic investigation of CREs and slows the search for additional non-coding pathological variants. As methods for single-cell transcriptomic and epigenomic profiling improve, it has become possible to identify cell-class-specific CREs in tens of thousands of individual cells simultaneously. The next challenge therefore will be to test the function of newly identified CREs in their native context of human retina-like cells. Human retinal organoids are a promising system in which to model the development and disorders of the human retina (Cowan et al., 2020; de Bruijn et al., 2020; Lu et al., 2020; Sridhar et al., 2020; Xie et al., 2020), but the degree of similarity in the cis-regulatory landscape between organoids and the retina was heretofore unexplored at the single-cell level. Ultimately, the work presented in this manuscript, identifying and comparing cell-class-specific CREs in the human retina and organoids, may help to elucidate the role of CREs in the development, function, and diseases of the retina.

Our approach was to take advantage of chromatin accessibility as one hallmark of CREs (Buenrostro et al., 2013; Thurman et al., 2012). We utilized snATAC-seq (Buenrostro et al., 2015; Cusanovich et al., 2015) to find candidate cell-class-specific CREs. Other methods are newly available to examine additional CRE features such as histone modifications and transcription factor binding at single-cell resolution (Kaya-Okur et al., 2019; Skene et al., 2018). However, for this study we reasoned that a strength of snATAC-seq is that it is relatively agnostic to the variety of CREs that it can identify and therefore can provide a broader inventory of candidate CREs to study. We then compared these data to snRNA-seq that we generated from the same tissues and timepoints and used recent software tools to statistically correlate individual CREs with candidate target genes, thereby addressing the significant outstanding issue of pairing CREs with their direct target genes, which can be >1 Mb away (Granja et al., 2021; Spielmann and Mundlos, 2016). Together these analyses and those presented in a companion paper (Lyu, 2021) provide a rich and accessible resource to investigate the role of CREs in retinal gene regulatory networks.

Future studies will need to manipulate CREs to confirm CRE-target gene associations and the biological functions of individual CREs. Currently, organoid and mouse models are two powerful and complementary systems to study retinal CRE function (Lyu, 2021; Miesfeld et al., 2020; Xie et al., 2020). Human organoids permit testing of CREs in the context of the intact human genome, while mouse models provide a physiologically relevant context with support cells and tissues like microglia, retinal vasculature, and retinal pigment epithelium. We therefore compared cell-class-specific CREs in the human retina to organoids and mice to determine which retinal CREs can be modeled in these systems. Moreover, we demonstrate a proof-of-concept functional investigation of a highly-conserved, disease-associated CRE in human retinal organoids by deleting the 5q14.3 enhancer. Our snATAC-seq analyses indicated that the dominant CRE at this human locus is enriched in retinal progenitor cells and Müller glia, which is consistent with the activity of its zebrafish homolog (Madelaine et al., 2018). We were able to confirm in a rigorous and quantitative manner that the functional target gene of this enhancer is the long non-coding RNA *LINC00461,* which serves as the primary transcript of miR-9-2. The evolutionarily conserved miR-9 family of microRNAs has a well-studied role in neural differentiation and has been investigated as one of several microRNAs involved in retinal progenitor timing and glial homeostasis (Coolen et al., 2013; La Torre et al., 2013; Wohl et al., 2017). Our findings gave us the opportunity to investigate the regulation and function of this individual family member (miR-9-2) in the disease-relevant human retinal organoid system. We found that regulation of miR-9-2 through this enhancer is required for the normal timing of retinal neurogenesis, but not for gliogenesis. We also found that this enhancer was essential for regulating gene-regulatory networks in mature Müller glia, providing insight into the association of this locus with the mature onset disorder MacTel, which is characterized by eventual loss of Müller glia and photoreceptor cells (Powner et al., 2013).

When analyzing dysregulated genes as a whole, we noted an interesting shift in metabolic pathway enrichment between enhancer KO progenitor cells and Müller glia indicating that e5q14.3 and miR-9-2 are playing distinct roles in these cell classes. The mTORC1 signaling, oxidative phosphorylation, hypoxia, and glycolysis pathways are all associated with upregulated gene expression in enhancer KO RPCs compared to controls. In contrast, these same genes are downregulated in KO Müller glia. While the upregulation of these pathways in KO progenitor cells may reflect the higher proliferative index in KO organoids, the downregulation of these pathways in Müller glia could have a long-term impact on the survival of these cells and on the metabolic support they provide to photoreceptors (Hurley et al., 2015; Shen et al., 2021). These findings could have significance to understanding 5q14.3-associated phenotypes including MacTel, which is known to be a metabolic disease linked to the dysregulation of serine synthesis through central carbon metabolism (Bonelli et al., 2020; Eade et al., 2021; Gantner et al., 2019). Furthermore, we also observe downregulated genes that are involved in vascular signaling, such as *FLT1, SEMA3A*, *COL18A1*, and *VEGFA,* in KO Müller glia. The miR-9 family has been implicated in vascular patterning (Madelaine et al., 2018; Madelaine et al., 2017), and Müller glia maintain connections with the surrounding vasculature (Vecino et al., 2016). These results also have important ramifications for the study of MacTel, which is characterized not only by the loss of Müller glia but also by aberrant retinal vascularization. We thus propose a model for e5q14.3 function within Müller glia, in which the enhancer directly targets *LINC00461* and regulates miR-9-2 expression, which suppresses target genes to maintain Müller glia homeostasis and promote regulation of retinal vasculature by Müller glia (Figure 7D).

While the current limitations of the organoid system prevent us from examining the role of this enhancer in the aged retina, where loss of Müller glia and vascular anomalies arise in MacTel, the results presented in this study may help to elucidate the very first molecular steps that lead to 5q14.3 associated pathology. Future experiments in mouse models may be able to address these outstanding questions. For now, it is our goal that the results presented here can serve as a roadmap for the next generation of studies of non-coding functional elements in the vertebrate retina and provide novel insights into the role of e5q14.3 and miR-9-2 in the developing and mature human retina.

## AUTHOR CONTRIBUTIONS

E.D.T., A.E.T., S.H.P., S.G., K.E., and T.J.C. designed and performed experiments and analyses associated with the manuscript. E.D.T. and A.E.T. guided bioinformatic analysis with A.E.T. serving as senior bioinformatician. P.L., T.H., J.Q., and S.B. generated mouse data and assisted with cross-species analyses. V.J. and M.B. assisted with human GWAS analyses. Study design was conceptualized by E.D.T., A.E.T, K.E., and T.J.C. E.D.T. and T.J.C. wrote the paper with input from all co-authors.

## ACKNOWLEDGEMENTS

We would like to thank all members of the Cherry Lab, James T. Bennett, the Lowy Medical Research Center, specifically Mari Gantner, Lea Scheppke, and Mike Dorrell as well as the LMRI Board of Scientific Governors for discussions about the project. Kimberly A. Aldinger and Thomas Vierbuchen provided independent feedback on this manuscript. We would also like to thank the Lowy family for their funding support of the MacTel Project, and Sang Sim, Megan Cook, Marti Lynn Moon and Jessica Orozoco for administrative assistance. This work was supported by grants from the NIH National Eye Institute R01EY028584 to T.J.C., R01EY020560 and U01EY027267 to S.B., R01EY029548 and P30EY001765 to J.Q., from The Brotman Baty Institute to T.J.C, an NIH T32 Grant Experimental Pathology of Cardiovascular Disease Training Grant HL007312 to E.D.T., and the Lowy Medical Research Institute to T.J.C. GWAS re-analysis was supported by the Australian Government National Health and Medical Research Council (NHMRC) Investigator Grant (1195236) to M.B. Additional funding to M.B. was provided by the Australian Independent Research Institute Infrastructure Support Scheme and the Victorian State Government Operational Infrastructure Program. The Birth Defects Research Laboratory at the University of Washington and Lion’s VisionGift of Portland, Oregon provided tissue samples. Lastly, we would like to acknowledge the tissue donors who made this study possible.

## SUPPLEMENTAL FIGURE LEGENDS

**Supplemental Figure 1: Mature retinal cell classes exhibit expected patterns of accessibility and gene expression over the course of their development.** (**A-G**) Changes in gene score values (Accessibility) and integrated gene expression values (Expression) over the course of pseudotime following trajectory analysis for each mature cell class. Accessibility values are plotted as log2(normalized counts +1), whereas Expression values are plotted as normalized counts. Warmer colors indicate higher pseudotime values.

**Supplemental Figure 2: Organoid cell classes exhibit expected patterns of accessibility and expression at maturity and during development.** (**A-B**) Feature plots showing the imputed gene score values (snATAC-seq, **A**) and the integrated gene expression values (snRNA-seq, **B**) for a variety of canonical retinal genes. Values are plotted as log2(normalized counts +1). Warmer colors indicate higher values. (**C-I**) Changes in gene score values (Accessibility) and integrated gene expression values (Expression) over the course of pseudotime following trajectory analysis for each mature cell class. Accessibility values are plotted as log2(normalized counts +1), whereas Expression values are plotted as normalized counts. Warmer colors indicate higher pseudotime values.

**Supplemental Figure 3: Enrichment of disease-gene-associated CREs between human retina, retinal organoids, and mouse retina.** (**A**) Heat map showing enrichment of CREs associated with known retinal disease genes within individual cell classes. Red sample text indicates organoid tissue, and black text indicates human tissue. (**B**) Heat map showing enrichment of CREs associated with known retinal disease genes within individual cell classes. Teal sample text indicates mouse tissue, and black text indicates human tissue.

**Supplemental Figure 4: Topologically associating domain (TAD) containing the 5q14.3 enhancer and candidate target genes.** Diagram showing the TAD housing e5q14.3 and the surrounding areas. (**A**) Genome-wide chromatin conformation capture data generated from hiPSC-derived retinal organoids described in de Bruijn et al., 2020. (**B**) Individual tracks showing CTCF binding, chromatin accessibility (ATAC-seq), permissive histone modifications (H3K27Ac), RNA-seq, and conservation across vertebrate species. The gray bar highlights the region in which e5q14.3 is located.

**Supplemental Figure 5: Analysis of differentially-expressed genes within RPCs.** (**A**) Volcano plot depicting all differentially expressed genes in KO primary RPCs. Dashed lines indicate significance thresholds: a significant p-value was set at p < 0.00001 (or -log(p) of 5) and significant log2 fold changes were set at ± 0.5. Gray dots indicate genes with no significant change. Blue dots indicate genes with a significant p-value but not a significant fold change. Red dots indicate genes with both a significant p-value and significant fold change. Green dots highlight genes that are potential e5q14.3 targets. Purple dots highlight genes that are predicted miR-9-2 targets. Green and purple dots are labeled with their gene names. (**B-C**) Gene set enrichment analysis of differentially-expressed genes in KO primary RPCs (**B**) and KO neurogenic RPCs (**C**). Normalized enrichment scores (NES) are plotted for all hallmark pathways. NES with an adjusted p-value of < 0.05 are shown in teal; NES with an adjusted p-value of ≥ 0.05 are shown in red. Pathways of interest are bolded.

**Supplemental Figure 6: Deletion of the 5q14.3 enhancer does not result in increased cell death.** (**A-B**) Maximum projections of sections from WT and KO organoids following TUNEL staining at week 16 (**A**) and week 20 (**B**). TUNEL-positive cells are shown in green, Chx10-positive cells (**A**) and Recoverin-positive cells (**B**) are shown in red, and DAPI-labeled nuclei are shown in blue. Scale bar = 25 µm (**A**) and 50 µm (**B**). (**A’-B’**) Quantification of TUNEL-positive cells in WT (gray) and KO (white) organoids at week 16 (**A’**) and week 20 (**B’**). n = 22 organoids (WT); n = 18 organoids (KO) (**A’**). n = 8 organoids (WT); n = 9 organoids (KO) (**B’**). Values are reported as mean ± SEM. Student’s unpaired t test. p = 0.5134 (**A’**); p = 0.7333 (**B’**).

**Supplemental Figure 7. Shift in fold-change in GSEA hallmark pathway genes between late RPCs and Müller glia.** (**A-D**) Plots of fold-changes of genes shared between late RPCs and Müller glia associated with the mTORC1 (**A**), Glycolysis (**B**), Hypoxia (**C**), and Notch (**D**) GSEA hallmark pathways. All genes have an adjusted p-value of < 0.01. Dashed line indicates a log2 fold change of 0.

## SUPPLEMENTAL TABLES

**Supplemental Table 1.** Table showing the number of nuclei (both that initially passed sequencing filter, and the final number following post-QC subsetting) as well as key quality control metrics for each sample.

**Supplemental Table 2.** Table of cell-class-specific marker genes used to assign cell class identity to different UMAP clusters.

**Supplemental Table 3.** Table depicting the normalized enrichment values of transcription factor binding sites in each cell class within the human retina snATAC-seq dataset. These values are visualized via heat map in Figure 1G.

**Supplemental Table 4.** Table of peak-to-gene correlation values for human retina, retinal organoid, and mouse retina snATAC-seq datasets.

**Supplemental Table 5.** List of retinal disease-associated genes used to filter the peak-to-gene list to identify disease-gene-associated CREs.

**Supplemental Table 6.** Table of differentially expressed genes in primary RPCs, neurogenic RPCs, and Müller glia in e5q14.3 KO retinal organoids.

## STAR METHODS

### RESOURCE AVAILABILITY

#### Lead Contact

Further information and requests for resources or reagents should be directed to the lead contact, Timothy J. Cherry.

#### Materials Availability

CRISPR plasmids that were used to generate the control and KO retinal organoids are available upon request.

#### Data and Code Availability

Data upload to GEO is in progress, and all unique code generated in this study will be made available through GitHub.

### EXPERIMENTAL MODEL AND SUBJECT DETAILS

#### Human Samples/Ethics Statement

The Seattle Children’s Hospital (SCH) Institutional Review Board reviewed and approved our procedures tissue procurement. Experiments were performed in accordance with SCH ethical and legal guidelines. Post-mortem adult human retinal tissues from de-identified donors were obtained from Lions VisionGift (Portland, OR, USA). The tissue was provided with de-identified medical records including time and cause of death, post-mortem interval prior to cryopreservation of tissue, age, perceived race, and sex. None of the donors had a prior medical history of ophthalmological conditions or interventions. Developing human retinas were obtained from the Birth Defects Research Laboratory at the University of Washington with ethics board approval and maternal written consent obtained before specimen collection. Information regarding sex and age accompanies the snATAC-seq and snRNA-seq files deposited with the gene expression omnibus (GEO: https://www.ncbi.nlm.nih.gov/geo/).

#### Human Pluripotent Cell Lines

The human induced pluripotent stem cell (hiPSC) lines (designated as cell line 1 and cell line 2) were derived from peripheral blood mononuclear cells from two separate, unaffected patients: one male donor, 61 years old; and one female donor, 58 years old. The male donor line was designated as cell line 1, and the female line was designated as cell line 2. These lines have been utilized and validated in previous studies (Eade et al., 2021; Gantner et al., 2019).

### METHOD DETAILS

#### Generation and Maintenance of hiPSCs

Reprogramming of hiPSC lines was performed by the Harvard iPS core facility using sendai virus for reprogramming factor delivery. The cell lines were verified for normal karyotypes (Cell Line Genetics Inc) and were contamination-free. hiPSCs were maintained on Matrigel (BD Biosciences) coated plates with mTeSR1 medium (STEMCELL Technologies). Cells were passaged every 3-4 days at approximately 80% confluency. Colonies containing clearly visible differentiated cells were marked and mechanically removed before passaging.

#### CRISPR Genome Editing

To generate the 1.2kbp excision at 5q14.3, we designed guide RNAs (gRNA) for a targeted double stranded breaks centered on either side of the rs17421627 SNP. The gRNA was designed using http://tools.genome-engineering.org and cloned it into the px330-puro-eGFP plasmid using Fast digest BpiI restriction enzyme (ThermoFisher). The gRNA sequences were:

gRNA 1: TAGCATATTAAGTGGTTGAC
gRNA 2: GCTAACCATATAAGTCGCAC

We transfected cell line 2 with the gRNA containing plasmid (1ug per gRNA) using the Neon Transfection System (ThermoFisher) with two 30sec 850V pulses. Transfected hiPSCs were selected using puromycin selection. Clones were established from single cell plating of puro selected hiPSCs and manually isolated to form clonal hiPSC lines following 10 days of expansion. We generated three independent homozygous excision clones and three wildtype control clones. The three control hiPSC lines were selected from independent hiPSC clones that underwent the same protocol without obtaining an excision event. Excisions were validated using Sanger sequencing.

Sanger sequencing primers:

Fwd 5’-3’- TCCTCTCTTCATAGCCTACC
Rev 5’-3’- CATTTCACACTACTCCCTGGA

#### Differentiation of Retinal Organoids

Retinal organoids were differentiated from hiPSCs between passage 10 and 20. Retinal organoids were differentiated as described in (https://pubmed.ncbi.nlm.nih.gov/33749682/). Timepoint 0 of differentiation refers to the formation of embryoid bodies. Following week 4 of differentiation, organoids were cultured in rotating cell suspension until week 18. Mature retinal organoids were cultured in Retinal Differentiation Media plus 10% FBS, 100 µM Taurine, and 2mM Glutamax starting at week 8 of differentiation.

#### Retinal Organoid Samples Used

All retinal organoids were derived from hiPSC cell line 2. For snATAC-seq experiments, 2 biological replicates were used per timepoint (5, 20, and 28 weeks). For snRNA-seq experiments, 2 biological replicates were used for 5-week control and KO organoids, and each replicate represented an independently edited sub-clone. For the other three timepoints in the snRNA-seq experiment (12, 20, and 28 weeks), three biological replicates were used per timepoint: two samples from one sub-clone, and the third from a different sub-clone. To account for variability between individual organoids, each biological replicate was comprised of a pool of 4-6 organoids

#### Nuclei Isolation

For snATAC-seq, nuclei were isolated following a modified version of 10x Genomics’ Nuclei Isolation from Mouse Brain Tissue for Single Cell ATAC Sequencing (CG000212, Rev. B), Protocol 2. 500 ul of chilled 0.1x Lysis Buffer (10 mMTris-HCl pH 7.4, 10 mM NaCl, 3 mM MgCl2, 0.01% Tween-20, 0.01% Nonidet P40 Substitute, 0.001% Digitonin, 1% BSA) was added to a tube of flash-frozen human retinal tissue or retinal organoid tissue and triturated 15-20 times. The suspension was then transferred to a dounce homogenizer cylinder and homogenized 10 times with an A pestle, then 10 times with a B pestle. The suspension was transferred to a new tube and incubated on ice for 5 minutes. The suspension was then pipette-mixed 10 times, then incubated on ice for another 10 minutes. 500 ul chilled Wash Buffer (10 mM Tris-HCl pH 7.4, 10 mM NaCl, 3 mM MgCl2, 1% BSA, 0.1% Tween-20) was then added, and the suspension was pipette-mixed 5 times and then passed through a 40 um Flowmi Cell Strainer into a new tube. Nuclei concentration was determined using a Countess II FL Automated Cell Counter, then the nuclei were centrifuged at 500 rcf for 5 minutes at 4 C. The supernatant was discarded and the nuclei were resuspended in the proper volume of chilled Diluted Nuclei Buffer (1x 10x Genomics Nuclei Buffer) to achieve a concentration range of 3,080-7,700 nuclei/ul, which is recommended for targeting 10,000 cells in the 10x Genomics snATAC-seq protocol. This recommended concentration was confirmed using the Countess II, and then the 10x snATACseq protocol was followed to completion.

For snRNA-seq, the same procedure was followed, but the Lysis and Wash buffers did not contain any Digitonin or Tween-20, and all buffers used also contained 0.2 U/ul RNase Inhibitor. For the final resuspension, the proper volume of chilled Diluted Nuclei Buffer was added to achieve a concentration range of 700-1200 nuclei/ul, again in order to target 10,000 cells for sequencing. This recommended concentration was confirmed using the Countess II, and then the 10x snRNA-seq protocol was followed to completion.

#### Sequencing

All sequencing was carried out by the Northwest Genomics Center at the University of Washington. Samples were sequenced on either a NovaSeq SP or NovaSeq S1 100-cycle flow cell, depending on how many samples were sequenced at a given time (SP for 4 or under, S1 for 5-9). All sequencing runs followed 10x Genomics’ recommended cycles and sequencing depth for both snRNA-seq and snATAC-seq. All data were returned in the FASTQ format.

#### Single Nucleus ATAC-seq Analysis

FASTQs were processed using the 10x Genomics cellranger-atac 1.2 pipeline. The count command was used to align reads to GRCh38/ hg38 and the fragment files were used as input for subsequent analysis. The resulting data were analyzed using the program suite ArchR (version 0.9.3). The output fragment files from cell ranger were used to generate Arrow files for each sample, doublets were identified, and all Arrow files were compiled into an ArchR project. After examining quality control metrics (transcriptional start site enrichment and the size distribution of unique nuclear fragments), doublets were removed from the project and an iterative LSI dimensionality reduction was performed. Following batch effect correction via Harmony, clustering and UMAP embedding were performed. Cluster-specific marker genes were identified by estimating gene expression via computation and imputation of gene scores. Next, snRNA-seq datasets for the same tissue types and timepoints (generated and analyzed via Seurat, see below) were integrated into the ArchR project using the unconstrained integration method, generating a Gene Integration Matrix linking actual gene expression from the scRNA-seq dataset with the inferred gene scores in the snATAC-seq dataset. Using both integrated gene expression and inferred gene scores in concert with *a priori* knowledge of cell-class-specific gene expression in the retina (see Table S2), we assigned cell class identities to the different clusters in the dataset. Peak calling was performed via MACS2, and cell-class-specific marker peaks were identified. Motif enrichment, motif footprinting, co-accessibility analysis, peak-to-gene linkage, and trajectory analysis were then performed.

#### Single Nucleus RNA-seq Analysis

FASTQs were processed using the 10x Genomics cellranger 3.1 pipeline. The count command was used to align reads to GRCh38/ hg38 and the counts matrices were used as input for subsequent analysis. The resulting data were analyzed using Seurat version 3.2.2. The output files from cell ranger were used to generate Seurat objects for each individual sample. Metadata corresponding to sample, timepoint, and genotype (when possible) were added, and then all objects were merged into one Seurat object. Quality control metrics were run (UMI number, gene number, and mitochondrial percentage), and the cells outside a specified range of feature counts and mitochondrial percentage were filtered out of the dataset (see code for details). In the case of the 5q14.3 enhancer knockout vs. control organoid dataset, the data then underwent normalization, feature selection, and scaling via Seurat’s SCTransform wrapper. Dimensionality reduction, UMAP embedding, and clustering were then performed. The object’s default assay was then switched to the RNA assay and renormalized. In the case of all other snRNA-seq datasets (human and non-knockout organoids), Seurat’s standard pipeline (not SCTransform) was used to perform normalization, feature selection, scaling, and dimensionality reduction. The data were then integrated using the Harmony package (version 0.1), after which UMAP embedding and clustering were performed. Differentially expressed genes within each cluster were then identified and used to determine the cell class identity of each cluster based on *a priori* knowledge of cell-class-specific gene expression (see Table S2). Clusters with abnormally low UMIs, high mitochondrial expression, or sample-specific origin were subset out of the dataset. Following subsetting, the object was re-clustered, differentially-expressed genes were re-identified, and final cell class identities were determined. To check for potential erosion of X-inactivation in hiPSC-derived cultures we initially examined expression of *XIST* on a per-sample basis and found variable levels of expression but no consistent trend based on timepoint or genotype. Upon further investigation we found no evidence of dysregulation of X chromosome transcripts in any samples, demonstrating that X inactivation was still intact. Cell class identities were assigned to clusters using the dplyr package. Differential expression, either across the entire dataset or on a cell-class-specific basis, was determined using the FindAllMarkers function with Idents set to genotype and logfc.threshold set to 0.

#### Correlation of Accessibility between Human and Organoid Datasets

To assess the relationship between samples and cell classes, we looked at the normalized coverage over a common set of peaks. If comparing between cell classes, barcodes of cells were retrieved from ArchR and the subset of reads were extracted BAM files by sinto. Peaks were called using MACS2 (with a q < 0.01 required). Peaks were subsequently merged and annotated which included quantification and normalization of reads around the peaks using HOMER. Spearman correlation was calculated within R and heat maps were plotted using the pheatmap package in R.

#### Correlation of Disease-Associated CREs between Human, Organoid, and Mouse Datasets

Within individual cell classes, peaks were called using MACS2 (q score < 0.000001) within ArchR. Peaks that had a Peak2GeneLinkage correlation (> 0.4) to a known retinal disease-associated gene (see Table S5) were kept. For pairwise comparisons bedtools intersect was used to find peaks that had more than 50% overlap between groups. For the human-mouse comparison, the UCSC Genome Browser was used to liftover peak coordinates from the mouse genome (mm10) to the human genome (hg38). Heat maps were plotted using the pheatmap package in R.

#### Generation of UCSC Browser Tracks

BAM files for individual cell classes were created using sinto. Bigwig files were generated using bamCoverage within the deepTools (3.4.3) suite of tools.

#### Volcano Plots

The Enhanced Volcano (version 1.4.0) package was used to generate volcano plots showing differentially expressed genes. The list of input genes and their associated fold changes and p-values was generated from Seurat’s FindAllMarkers function. Only the expression changes in KO organoids were used as an input. The gene *XIST* was removed from the datasets prior to volcano plot input to represent the dynamic range of other transcripts, but is still present in Table S6. All adjusted p-values that were listed as 0 were set to 2.23E-308, the smallest non-zero value R can compute, to avoid undefined values. The significant p-value threshold was set at p = 0.000001, and the significant log2 fold change was set at ±0.5.

#### Gene Set Enrichment Analysis

Cell-class-specific differential expression was generated using the presto package (version 1.0.0). The expression changes of all genes, regardless of adjusted p-value, within KO organoids were listed in rank order by their area under the curve (AUC) statistic. This ranked list was then input to the fgsea package (version 1.16.0) using the Hallmark gene set list from MSigDB. Normalized enrichment scores were then plotted, with a significance threshold set at p ≤ 0.05.

#### Immunohistochemistry

*Fixation and cryosectioning*: Organoid tissue was fixed in 4% PFA in PBS for 10 mins, washed in PBS, and then submerged in 20% sucrose in PBS overnight. Tissues were embedded in O.C.T compound and frozen at −80°C. Cryosectioning was done at 14 μm and sections were mounted on glass polylysine coated slides. *Antibody staining*: Prior to primary antibody addition, sections were blocked with 5% donkey serum in PBS with 0.1% tween. Primary antibodies were added to samples at 4°C and incubated overnight. Following primary antibody incubation, sections were washed 3 x 10 mins in PBS. Sections were incubated with species-specific secondary antibodies (1:1000 dilution) for 2 hours. Following secondary antibody incubation, samples were washed 3 x 10 mins in PBS and mounted in mounting media.

*Primary antibody working dilutions*: rabbit anti-Recoverin (1:500); mouse anti-RLBP (1:200); rabbit anti-SOX9 (1:200); sheep anti-CHX10 (1:500).

#### Proliferation Assay

EdU birth dating was performed using the EdU-AlexaFluor 488 kit (Invitrogen C10337). The protocol was modified to accommodate cryosections on glass slides by using a wax pen to create a dam to hold solutions in place. EdU staining protocol was performed following cryosectioning on to glass slides. Following EdU staining sections underwent the standard immunostaining protocol described above.

#### Quantification of Cell Death

TUNEL staining was performed using In Situ Cell Death Detection Fluorescein kit (Sigma cat# 11684795910) prior to addition of primary antibody. Overlapping TUNEL positive staining and DAPI staining within a recoverin positive cell was counted as cell death within a photoreceptor. Cell death was normalized to the area of CHX10 (week 16) or Recoverin (week 20) staining in the retinal organoid.

### QUANTIFICATION AND STATISTICAL ANALYSES

Statistical analyses not performed in R via ArchR or Seurat were performed in GraphPad Prism 6.0. Student’s unpaired t test was used for comparisons of two groups, whereas a two-way ANOVA with a Sidak’s multiple comparisons test was used for comparisons of three or more groups. Statistical significance was set at p ≤ 0.05.

### KEY RESOURCES TABLE

| **REAGENT or RESOURCE** | **SOURCE** | **IDENTIFIER** |
| --- | --- | --- |
| **Antibodies** |  |  |
| Rabbit polyclonal anti-Recoverin | Millipore | Cat# AB5585 |
| Sheep polyclonal anti-Chx10 | Millipore | Cat# AB9016 |
| Mouse monoclonal anti-RLBP1 | Abcam | Cat# ab15051 |
| Rabbit polyclonal anti-Sox9 | Millipore | Cat# AB5535 |
| Alexa Fluor 555 Highly Cross-Adsorbed Donkey anti-Mouse IgG (H+L) | Invitrogen | Cat# A-21202 |
| Alexa Fluor 555 Highly Cross-Adsorbed Donkey anti-Rabbit IgG (H+L) | Invitrogen | Cat# A-31572 |
| Alexa Fluor 555 Highly Cross-Adsorbed Donkey anti-Mouse IgG (H+L) | Invitrogen | Cat# A-31570 |
| Alexa Fluor 555 Highly Cross-Adsorbed Donkey anti-Sheep IgG (H+L) | Invitrogen | Cat# A-21436 |
| **Biological samples** |  |  |
| Developing human retinal tissue | Birth Defects Research Laboratory, University of Washington, Seattle, WA |  |
| Adult human retinal tissue | Lions VisionGift, Portland, OR |  |
| Retinal organoids | Lowy Medical Research Institute | Generated from hiPSCs listed below |
| **Chemicals, peptides, and recombinant proteins** |  |  |
| Click-iT EdU Cell Proliferation Kit, Alexa Fluor 488 | Invitrogen | Cat# C10337 |
| In Situ Cell Death Detection Kit, Fluorescein | Sigma | Cat# 11684795910 |
| 5% Digitonin | Invitrogen | Cat# BN2006 |
| Sigma Protector RNase Inhibitor | Sigma-Aldrich | Cat# 3335399001 |
| SPRIselect Reagent Kit | Beckman Coulter | Cat# B23317 |
| Low TE Buffer | Invitrogen | Cat# 12090-105 |
| 10% Tween-20 | Bio-Rad | Cat# 166-2404 |
| mTeSR-1 | StemCell Technologies | Cat# 85850 |
| Matrigel | Corning | Cat# 354230 |
| Y-27632 dihydrochloride | Sigma | Cat# Y0503 |
| DMEM | Gibco | Cat# 11995065 |
| DMEM/F12 | Gibco | Cat# 11330057 |
| N2 Supplement | Gibco | Cat# 17502-048 |
| NEAA | Gibco | Cat# 11140-050 |
| Pen-Strep | Gibco | Cat# 15140-122 |
| Heparin | StemCell Technologies | Cat# 7980 |
| B27 Supplement | Gibco | Cat# 12587-010 |
| Glutamax | Gibco | Cat# 35050-061 |
| Taurine | Sigma | Cat# T0625 |
| Fetal Bovine Serum | Corning | Cat# 45001-108 |
| **Critical commercial assays** |  |  |
| Chromium Next GEM Single Cell 3' GEM, Library & Gel Bead Kit v3.1 | 10x Genomics | Cat# 1000128 |
| Chromium Next GEM Single Cell ATAC Library & Gel Bead Kit v1.1 | 10x Genomics | Cat# 1000176 |
| Agilent High Sensitivity DNA Kit | Agilent | Cat# 5067-4626 |
| Qubit dsDNA HS Assay Kit | Invitrogen | Cat# Q32851 |
| **Deposited data** |  |  |
| snATAC-seq data | This paper | GEO access in progress |
| snRNA-seq data | This paper | GEO access in progress |
| UCSC Browser Tracks | This paper | URL pending |
| **Experimental models: Cell lines** |  |  |
| hiPSC cell line used to generate organoids | Lowy Medical Research Institute | Available upon request |
| **Software and algorithms** |  |  |
| Cell Ranger 3.1 | 10x Genomics | https://www.10xgenomics.com/ |
| Cell Ranger ATAC 1.2 | 10x Genomics | https://www.10xgenomics.com/ |
| Seurat v 3.2.2 | (Stuart et al., 2019) | https://satijalab.org/seurat/ |
| ArchR v 0.9.5 | (Granja et al., 2021) | https://www.archrproject.com/index.html |
| Harmony v 1.0 | (Korsunsky et al., 2019) | https://portals.broadinstitute.org/harmony/ |
| Presto v 1.0.0 | Immunogenomics | https://github.com/immunogenomics/presto |
| dplyr v 1.0.4 | Hadley Wickham | https://github.com/tidyverse/dplyr |
| Enhanced Volcano v 1.4.0 | Blighe, Rana, and Lewis (2018) | https://github.com/kevinblighe/EnhancedVolcano |
| MACS v 2.2.7.1 | (Zhang et al., 2008) | https://github.com/macs3-project/MACS |
| HOMER v 4.9.1 | (Heinz et al., 2010) | http://homer.ucsd.edu/homer/ |
| pheatmap v 1.0.12 | Raivo Kolde | https://cran.r-project.org/web/packages/pheatmap/index.html |
| fgsea v 1.16.0 | Korotkevich et al., 2019 | https://github.com/ctlab/fgsea |
| sinto v 0.7.1 | Tim Stuart, Warren W. Kretzschmar | https://github.com/timoast/sinto |
| deepTools v 3.4.3 | (Ramirez et al., 2016) | https://github.com/deeptools/deepTools/blob/develop/docs/index.rst |
| R v 3.6.3, v 4.0.4 | The R Project | https://www.r-project.org/ |
| GraphPad Prism 6.0 | GraphPad Software | https://www.graphpad.com/scientific-software/prism/ |
| FIJI | (Schindelin et al., 2012) | https://imagej.net/software/fiji/ |
| Photoshop | Adobe | https://www.adobe.com/products/photoshop.html |
| BioRender | BioRender | https://biorender.com/ |
| TargetScan | (Agarwal et al., 2015) | http://www.targetscan.org/vert_72/ |

## REFERENCES

Agarwal, V., Bell, G.W., Nam, J.W., and Bartel, D.P. (2015). Predicting effective microRNA target sites in mammalian mRNAs. Elife 4.

Aldiri, I., Xu, B., Wang, L., Chen, X., Hiler, D., Griffiths, L., Valentine, M., Shirinifard, A., Thiagarajan, S., Sablauer, A., et al. (2017). The Dynamic Epigenetic Landscape of the Retina During Development, Reprogramming, and Tumorigenesis. Neuron 94, 550–568 e510.

Andzelm, M.M., Cherry, T.J., Harmin, D.A., Boeke, A.C., Lee, C., Hemberg, M., Pawlyk, B., Malik, A.N., Flavell, S.W., Sandberg, M.A., et al. (2015). MEF2D drives photoreceptor development through a genome-wide competition for tissue-specific enhancers. Neuron 86, 247–263.

Bhatia, S., Bengani, H., Fish, M., Brown, A., Divizia, M.T., de Marco, R., Damante, G., Grainger, R., van Heyningen, V., and Kleinjan, D.A. (2013). Disruption of autoregulatory feedback by a mutation in a remote, ultraconserved PAX6 enhancer causes aniridia. Am J Hum Genet 93, 1126–1134.

Bonelli, R., Jackson, V.E., Prasad, A., Munro, J.E., Farashi, S., Heeren, T.F.C., Pontikos, N., Scheppke, L., Friedlander, M., MacTel, C., et al. (2021). Identification of genetic factors influencing metabolic dysregulation and retinal support for MacTel, a retinal disorder. Commun Biol 4, 274.

Bonelli, R., Woods, S.M., Ansell, B.R.E., Heeren, T.F.C., Egan, C.A., Khan, K.N., Guymer, R., Trombley, J., Friedlander, M., Bahlo, M., et al. (2020). Systemic lipid dysregulation is a risk factor for macular neurodegenerative disease. Sci Rep 10, 12165.

Brodie-Kommit, J., Clark, B.S., Shi, Q., Shiau, F., Kim, D.W., Langel, J., Sheely, C., Ruzycki, P.A., Fries, M., Javed, A., et al. (2021). Atoh7-independent specification of retinal ganglion cell identity. Sci Adv 7.

Buenrostro, J.D., Giresi, P.G., Zaba, L.C., Chang, H.Y., and Greenleaf, W.J. (2013). Transposition of native chromatin for fast and sensitive epigenomic profiling of open chromatin, DNA-binding proteins and nucleosome position. Nat Methods 10, 1213–1218.

Buenrostro, J.D., Wu, B., Litzenburger, U.M., Ruff, D., Gonzales, M.L., Snyder, M.P., Chang, H.Y., and Greenleaf, W.J. (2015). Single-cell chromatin accessibility reveals principles of regulatory variation. Nature 523, 486–490.

Capowski, E.E., Samimi, K., Mayerl, S.J., Phillips, M.J., Pinilla, I., Howden, S.E., Saha, J., Jansen, A.D., Edwards, K.L., Jager, L.D., et al. (2019). Reproducibility and staging of 3D human retinal organoids across multiple pluripotent stem cell lines. Development 146.

Chan, C.S.Y., Lonfat, N., Zhao, R., Davis, A.E., Li, L., Wu, M.R., Lin, C.H., Ji, Z., Cepko, C.L., and Wang, S. (2020). Cell type- and stage-specific expression of Otx2 is regulated by multiple transcription factors and cis-regulatory modules in the retina. Development 147.

Cherry, T.J., Yang, M.G., Harmin, D.A., Tao, P., Timms, A.E., Bauwens, M., Allikmets, R., Jones, E.M., Chen, R., De Baere, E., et al. (2020). Mapping the cis-regulatory architecture of the human retina reveals noncoding genetic variation in disease. Proc Natl Acad Sci U S A 117, 9001–9012.

Clark, B.S., Stein-O’Brien, G.L., Shiau, F., Cannon, G.H., Davis-Marcisak, E., Sherman, T., Santiago, C.P., Hoang, T.V., Rajaii, F., James-Esposito, R.E., et al. (2019). Single-Cell RNA-Seq Analysis of Retinal Development Identifies NFI Factors as Regulating Mitotic Exit and Late-Born Cell Specification. Neuron 102, 1111–1126 e1115.

Coolen, M., Katz, S., and Bally-Cuif, L. (2013). miR-9: a versatile regulator of neurogenesis. Front Cell Neurosci 7, 220.

Cowan, C.S., Renner, M., De Gennaro, M., Gross-Scherf, B., Goldblum, D., Hou, Y., Munz, M., Rodrigues, T.M., Krol, J., Szikra, T., et al. (2020). Cell Types of the Human Retina and Its Organoids at Single-Cell Resolution. Cell 182, 1623–1640 e1634.

Cusanovich, D.A., Daza, R., Adey, A., Pliner, H.A., Christiansen, L., Gunderson, K.L., Steemers, F.J., Trapnell, C., and Shendure, J. (2015). Multiplex single cell profiling of chromatin accessibility by combinatorial cellular indexing. Science 348, 910–914.

de Bruijn, S.E., Fiorentino, A., Ottaviani, D., Fanucchi, S., Melo, U.S., Corral-Serrano, J.C., Mulders, T., Georgiou, M., Rivolta, C., Pontikos, N., et al. (2020). Structural Variants Create New Topological-Associated Domains and Ectopic Retinal Enhancer-Gene Contact in Dominant Retinitis Pigmentosa. Am J Hum Genet 107, 802–814.

Eade, K., Gantner, M.L., Hostyk, J.A., Nagasaki, T., Giles, S., Fallon, R., Harkins-Perry, S., Baldini, M., Lim, E.W., Scheppke, L., et al. (2021). Serine biosynthesis defect due to haploinsufficiency of PHGDH causes retinal disease. Nat Metab 3, 366–377.

Eiraku, M., Takata, N., Ishibashi, H., Kawada, M., Sakakura, E., Okuda, S., Sekiguchi, K., Adachi, T., and Sasai, Y. (2011). Self-organizing optic-cup morphogenesis in three-dimensional culture. Nature 472, 51–56.

Erkman, L., McEvilly, R.J., Luo, L., Ryan, A.K., Hooshmand, F., O’Connell, S.M., Keithley, E.M., Rapaport, D.H., Ryan, A.F., and Rosenfeld, M.G. (1996). Role of transcription factors Brn-3.1 and Brn-3.2 in auditory and visual system development. Nature 381, 603–606.

Fritsche, L.G., Igl, W., Bailey, J.N., Grassmann, F., Sengupta, S., Bragg-Gresham, J.L., Burdon, K.P., Hebbring, S.J., Wen, C., Gorski, M., et al. (2016). A large genome-wide association study of age-related macular degeneration highlights contributions of rare and common variants. Nat Genet 48, 134–143.

Furukawa, T., Morrow, E.M., and Cepko, C.L. (1997). Crx, a novel otx-like homeobox gene, shows photoreceptor-specific expression and regulates photoreceptor differentiation. Cell 91, 531–541.

Gachon, F., Fonjallaz, P., Damiola, F., Gos, P., Kodama, T., Zakany, J., Duboule, D., Petit, B., Tafti, M., and Schibler, U. (2004). The loss of circadian PAR bZip transcription factors results in epilepsy. Genes Dev 18, 1397–1412.

Gan, L., Xiang, M., Zhou, L., Wagner, D.S., Klein, W.H., and Nathans, J. (1996). POU domain factor Brn-3b is required for the development of a large set of retinal ganglion cells. Proc Natl Acad Sci U S A 93, 3920–3925.

Gantner, M.L., Eade, K., Wallace, M., Handzlik, M.K., Fallon, R., Trombley, J., Bonelli, R., Giles, S., Harkins-Perry, S., Heeren, T.F.C., et al. (2019). Serine and Lipid Metabolism in Macular Disease and Peripheral Neuropathy. N Engl J Med 381, 1422–1433.

Gao, X.R., Huang, H., and Kim, H. (2019). Genome-wide association analyses identify 139 loci associated with macular thickness in the UK Biobank cohort. Hum Mol Genet 28, 1162–1172.

Ghiasvand, N.M., Rudolph, D.D., Mashayekhi, M., Brzezinski, J.A.t., Goldman, D., and Glaser, T. (2011). Deletion of a remote enhancer near ATOH7 disrupts retinal neurogenesis, causing NCRNA disease. Nat Neurosci 14, 578–586.

Goodson, N.B., Kaufman, M.A., Park, K.U., and Brzezinski, J.A.t. (2020). Simultaneous deletion of Prdm1 and Vsx2 enhancers in the retina alters photoreceptor and bipolar cell fate specification, yet differs from deleting both genes. Development 147.

Granja, J.M., Corces, M.R., Pierce, S.E., Bagdatli, S.T., Choudhry, H., Chang, H.Y., and Greenleaf, W.J. (2021). ArchR is a scalable software package for integrative single-cell chromatin accessibility analysis. Nat Genet 53, 403–411.

Han, X., Gharahkhani, P., Mitchell, P., Liew, G., Hewitt, A.W., and MacGregor, S. (2020). Genome-wide meta-analysis identifies novel loci associated with age-related macular degeneration. J Hum Genet 65, 657–665.

Heinz, S., Benner, C., Spann, N., Bertolino, E., Lin, Y.C., Laslo, P., Cheng, J.X., Murre, C., Singh, H., and Glass, C.K. (2010). Simple combinations of lineage-determining transcription factors prime cis-regulatory elements required for macrophage and B cell identities. Mol Cell 38, 576–589.

Hoshino, A., Ratnapriya, R., Brooks, M.J., Chaitankar, V., Wilken, M.S., Zhang, C., Starostik, M.R., Gieser, L., La Torre, A., Nishio, M., et al. (2017). Molecular Anatomy of the Developing Human Retina. Dev Cell 43, 763–779 e764.

Hurley, J.B., Lindsay, K.J., and Du, J. (2015). Glucose, lactate, and shuttling of metabolites in vertebrate retinas. J Neurosci Res 93, 1079–1092.

Ikram, M.K., Sim, X., Jensen, R.A., Cotch, M.F., Hewitt, A.W., Ikram, M.A., Wang, J.J., Klein, R., Klein, B.E., Breteler, M.M., et al. (2010). Four novel Loci (19q13, 6q24, 12q24, and 5q14) influence the microcirculation in vivo. PLoS Genet 6, e1001184.

Jadhav, A.P., Cho, S.H., and Cepko, C.L. (2006a). Notch activity permits retinal cells to progress through multiple progenitor states and acquire a stem cell property. Proc Natl Acad Sci U S A 103, 18998–19003.

Jadhav, A.P., Mason, H.A., and Cepko, C.L. (2006b). Notch 1 inhibits photoreceptor production in the developing mammalian retina. Development 133, 913–923.

Jin, K., Jiang, H., Xiao, D., Zou, M., Zhu, J., and Xiang, M. (2015). Tfap2a and 2b act downstream of Ptf1a to promote amacrine cell differentiation during retinogenesis. Mol Brain 8, 28.

Kaya-Okur, H.S., Wu, S.J., Codomo, C.A., Pledger, E.S., Bryson, T.D., Henikoff, J.G., Ahmad, K., and Henikoff, S. (2019). CUT&Tag for efficient epigenomic profiling of small samples and single cells. Nat Commun 10, 1930.

Korsunsky, I., Millard, N., Fan, J., Slowikowski, K., Zhang, F., Wei, K., Baglaenko, Y., Brenner, M., Loh, P.R., and Raychaudhuri, S. (2019). Fast, sensitive and accurate integration of single-cell data with Harmony. Nat Methods 16, 1289–1296.

La Torre, A., Georgi, S., and Reh, T.A. (2013). Conserved microRNA pathway regulates developmental timing of retinal neurogenesis. Proc Natl Acad Sci U S A 110, E2362–2370.

Liao, S.M., Zheng, W., Zhu, J., Lewis, C.A., Delgado, O., Crowley, M.A., Buchanan, N.M., Jaffee, B.D., and Dryja, T.P. (2017). Specific correlation between the major chromosome 10q26 haplotype conferring risk for age-related macular degeneration and the expression of HTRA1. Mol Vis 23, 318–333.

Lu, Y., Shiau, F., Yi, W., Lu, S., Wu, Q., Pearson, J.D., Kallman, A., Zhong, S., Hoang, T., Zuo, Z., et al. (2020). Single-Cell Analysis of Human Retina Identifies Evolutionarily Conserved and Species-Specific Mechanisms Controlling Development. Dev Cell 53, 473–491 e479.

Lyu, P.H., T.; Santiago, C.P.; Thomas, E.D.; Timms, A.E.; Appel, H.; Gimmen, M.; Nguyet, L.; Jiang, L.; Kim, D.W.; Chen, S.; Espinoza, D.; Telger, A.E.; Weir, K.; Clark, B.S.; Cherry, T.J.; Qian, J.; and Blackshaw, S. (2021). Integrated multiomic analysis identifies gene regulatory networks controlling temporal patterning, neurogenesis and cell fate specification in the mammalian retina. bioRxiv 2021/454200.

Madelaine, R., Notwell, J.H., Skariah, G., Halluin, C., Chen, C.C., Bejerano, G., and Mourrain, P. (2018). A screen for deeply conserved non-coding GWAS SNPs uncovers a MIR-9-2 functional mutation associated to retinal vasculature defects in human. Nucleic Acids Res 46, 3517–3531.

Madelaine, R., Sloan, S.A., Huber, N., Notwell, J.H., Leung, L.C., Skariah, G., Halluin, C., Pasca, S.P., Bejerano, G., Krasnow, M.A., et al. (2017). MicroRNA-9 Couples Brain Neurogenesis and Angiogenesis. Cell Rep 20, 1533–1542.

Maurano, M.T., Humbert, R., Rynes, E., Thurman, R.E., Haugen, E., Wang, H., Reynolds, A.P., Sandstrom, R., Qu, H., Brody, J., et al. (2012). Systematic localization of common disease-associated variation in regulatory DNA. Science 337, 1190–1195.

Miesfeld, J.B., Ghiasvand, N.M., Marsh-Armstrong, B., Marsh-Armstrong, N., Miller, E.B., Zhang, P., Manna, S.K., Zawadzki, R.J., Brown, N.L., and Glaser, T. (2020). The Atoh7 remote enhancer provides transcriptional robustness during retinal ganglion cell development. Proc Natl Acad Sci U S A 117, 21690–21700.

Nathans, J., Davenport, C.M., Maumenee, I.H., Lewis, R.A., Hejtmancik, J.F., Litt, M., Lovrien, E., Weleber, R., Bachynski, B., Zwas, F., et al. (1989). Molecular genetics of human blue cone monochromacy. Science 245, 831–838.

Norrie, J.L., Lupo, M.S., Xu, B., Al Diri, I., Valentine, M., Putnam, D., Griffiths, L., Zhang, J., Johnson, D., Easton, J., et al. (2019). Nucleome Dynamics during Retinal Development. Neuron 104, 512–528 e511.

Oliver, P.L., Chodroff, R.A., Gosal, A., Edwards, B., Cheung, A.F., Gomez-Rodriguez, J., Elliot, G., Garrett, L.J., Lickiss, T., Szele, F., et al. (2015). Disruption of Visc-2, a Brain-Expressed Conserved Long Noncoding RNA, Does Not Elicit an Overt Anatomical or Behavioral Phenotype. Cereb Cortex 25, 3572–3585.

Powner, M.B., Gillies, M.C., Zhu, M., Vevis, K., Hunyor, A.P., and Fruttiger, M. (2013). Loss of Muller’s cells and photoreceptors in macular telangiectasia type 2. Ophthalmology 120, 2344–2352.

Ramirez, F., Ryan, D.P., Gruning, B., Bhardwaj, V., Kilpert, F., Richter, A.S., Heyne, S., Dundar, F., and Manke, T. (2016). deepTools2: a next generation web server for deep-sequencing data analysis. Nucleic Acids Res 44, W160–165.

Scerri, T.S., Quaglieri, A., Cai, C., Zernant, J., Matsunami, N., Baird, L., Scheppke, L., Bonelli, R., Yannuzzi, L.A., Friedlander, M., et al. (2017). Genome-wide analyses identify common variants associated with macular telangiectasia type 2. Nat Genet 49, 559–567.

Schindelin, J., Arganda-Carreras, I., Frise, E., Kaynig, V., Longair, M., Pietzsch, T., Preibisch, S., Rueden, C., Saalfeld, S., Schmid, B., et al. (2012). Fiji: an open-source platform for biological-image analysis. Nat Methods 9, 676–682.

Shen, W., Lee, S.R., Mathai, A.E., Zhang, R., Du, J., Yam, M.X., Pye, V., Barnett, N.L., Rayner, C.L., Zhu, L., et al. (2021). Effect of selectively knocking down key metabolic genes in Muller glia on photoreceptor health. Glia 69, 1966–1986.

Skene, P.J., Henikoff, J.G., and Henikoff, S. (2018). Targeted in situ genome-wide profiling with high efficiency for low cell numbers. Nat Protoc 13, 1006–1019.

Spielmann, M., and Mundlos, S. (2016). Looking beyond the genes: the role of non-coding variants in human disease. Hum Mol Genet 25, R157–R165.

Sridhar, A., Hoshino, A., Finkbeiner, C.R., Chitsazan, A., Dai, L., Haugan, A.K., Eschenbacher, K.M., Jackson, D.L., Trapnell, C., Bermingham-McDonogh, O., et al. (2020). Single-Cell Transcriptomic Comparison of Human Fetal Retina, hPSC-Derived Retinal Organoids, and Long-Term Retinal Cultures. Cell Rep 30, 1644–1659 e1644.

Stuart, T., Butler, A., Hoffman, P., Hafemeister, C., Papalexi, E., Mauck, W.M., 3rd, Hao, Y., Stoeckius, M., Smibert, P., and Satija, R. (2019). Comprehensive Integration of Single-Cell Data. Cell 177, 1888–1902 e1821.

Thurman, R.E., Rynes, E., Humbert, R., Vierstra, J., Maurano, M.T., Haugen, E., Sheffield, N.C., Stergachis, A.B., Wang, H., Vernot, B., et al. (2012). The accessible chromatin landscape of the human genome. Nature 489, 75–82.

Vecino, E., Rodriguez, F.D., Ruzafa, N., Pereiro, X., and Sharma, S.C. (2016). Glia-neuron interactions in the mammalian retina. Prog Retin Eye Res 51, 1–40.

Wang, S., Sengel, C., Emerson, M.M., and Cepko, C.L. (2014). A gene regulatory network controls the binary fate decision of rod and bipolar cells in the vertebrate retina. Dev Cell 30, 513–527.

Wohl, S.G., Jorstad, N.L., Levine, E.M., and Reh, T.A. (2017). Muller glial microRNAs are required for the maintenance of glial homeostasis and retinal architecture. Nat Commun 8, 1603.

Wu, F., Sapkota, D., Li, R., and Mu, X. (2012). Onecut 1 and Onecut 2 are potential regulators of mouse retinal development. J Comp Neurol 520, 952–969.

Xie, H., Zhang, W., Zhang, M., Akhtar, T., Li, Y., Yi, W., Sun, X., Zuo, Z., Wei, M., Fang, X., et al. (2020). Chromatin accessibility analysis reveals regulatory dynamics of developing human retina and hiPSC-derived retinal organoids. Sci Adv 6, eaay5247.

Zhang, Y., Liu, T., Meyer, C.A., Eeckhoute, J., Johnson, D.S., Bernstein, B.E., Nusbaum, C., Myers, R.M., Brown, M., Li, W., et al. (2008). Model-based analysis of ChIP-Seq (MACS). Genome Biol 9, R137.

