## Supplementary figures and images for "Multi-omic Analysis of Developing Human Retina and Organoids Reveals Cell-Specific Cis-Regulatory Elements and Mechanisms of Non-Coding Genetic Disease Risk"

### Figure S1

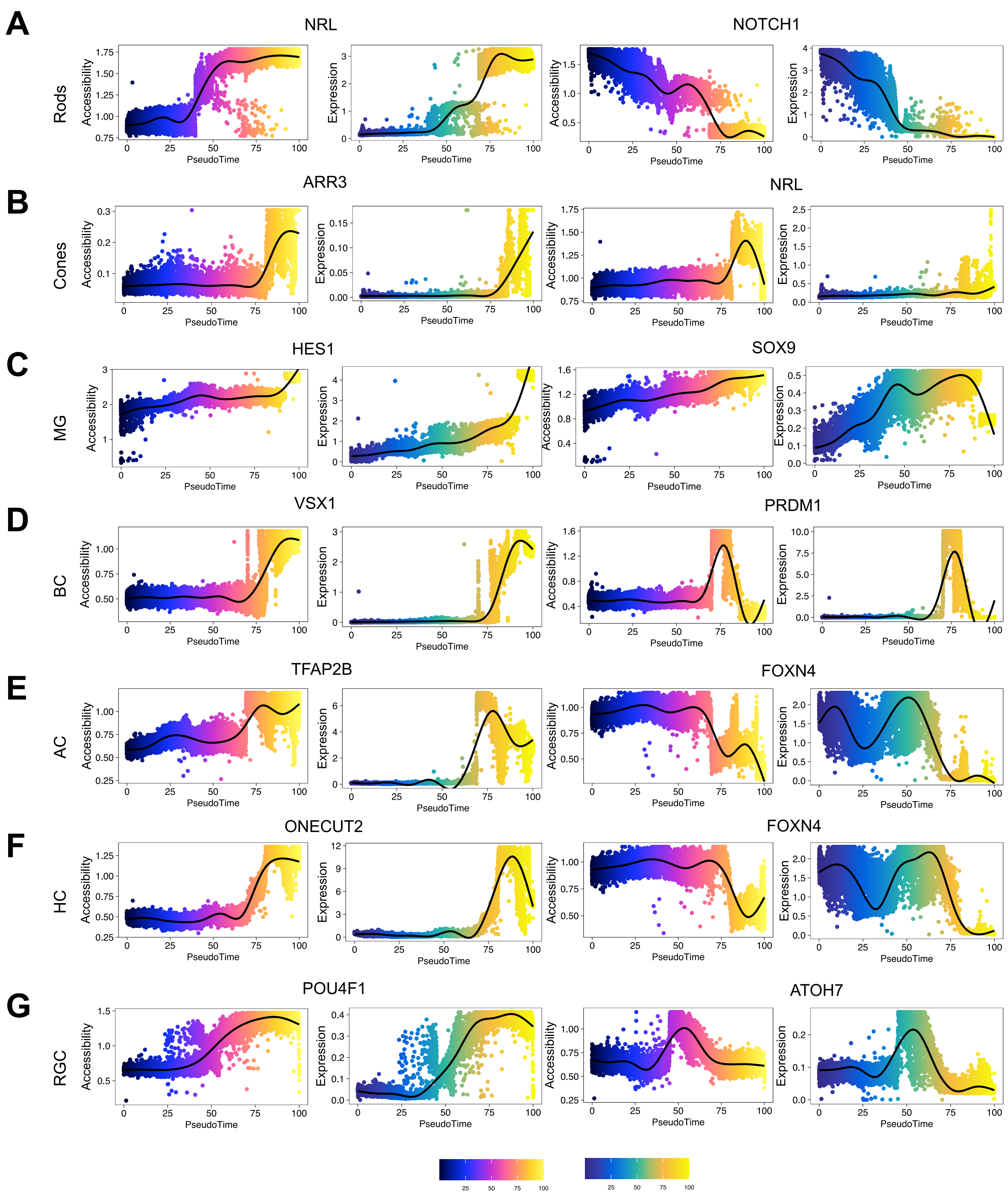

### Figure S2

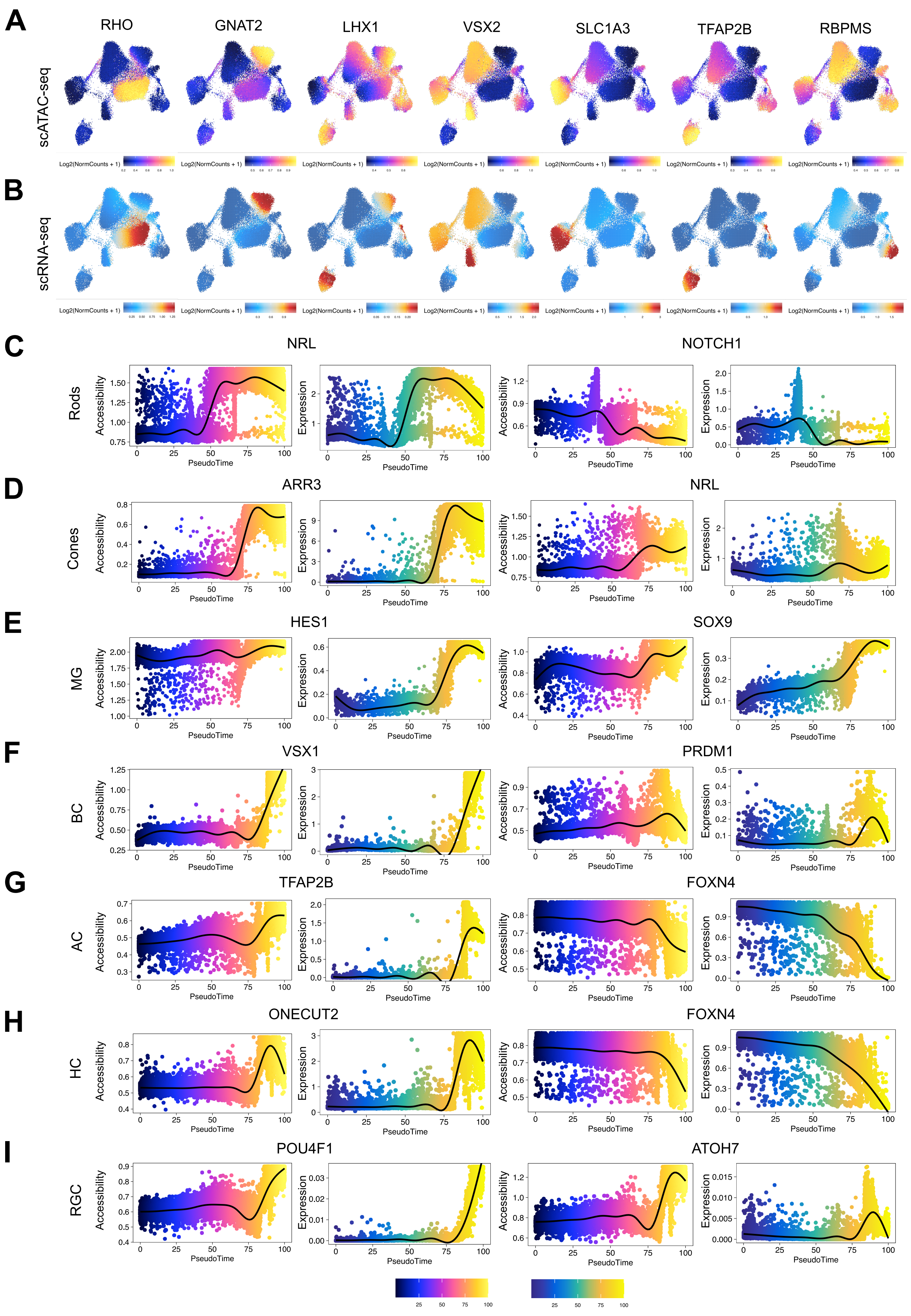

### Figure S4

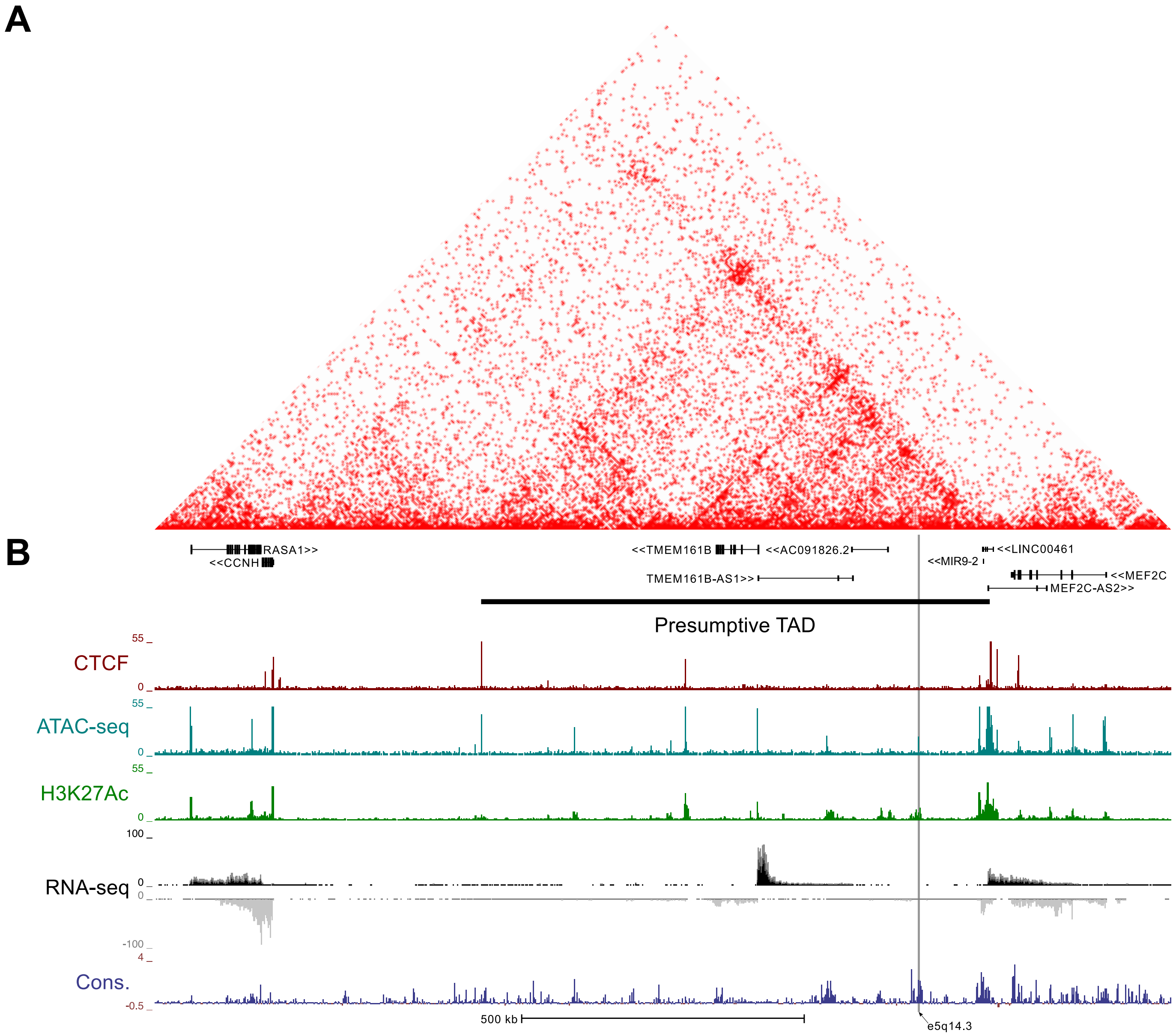

### Figure S5

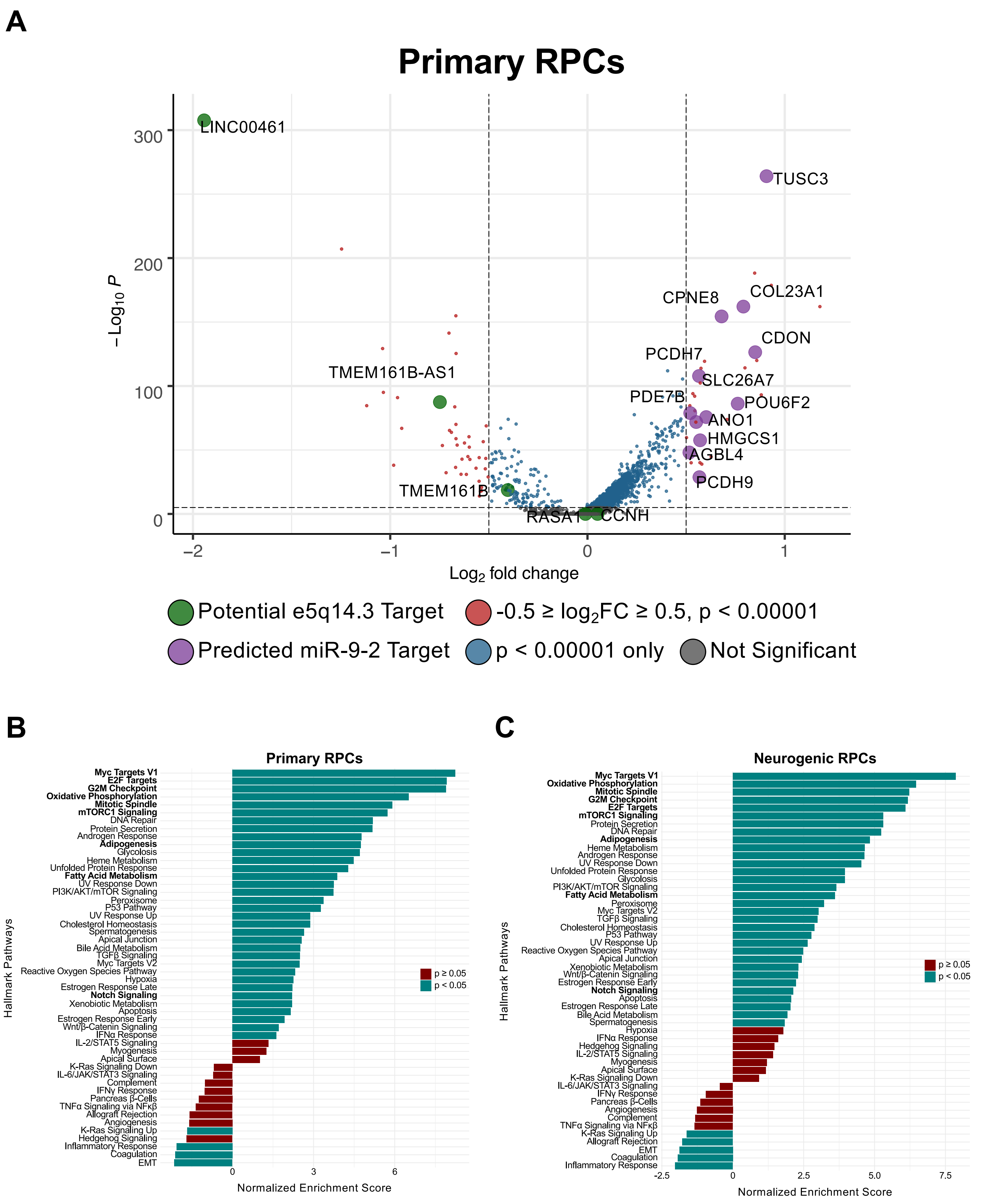

### Figure S6

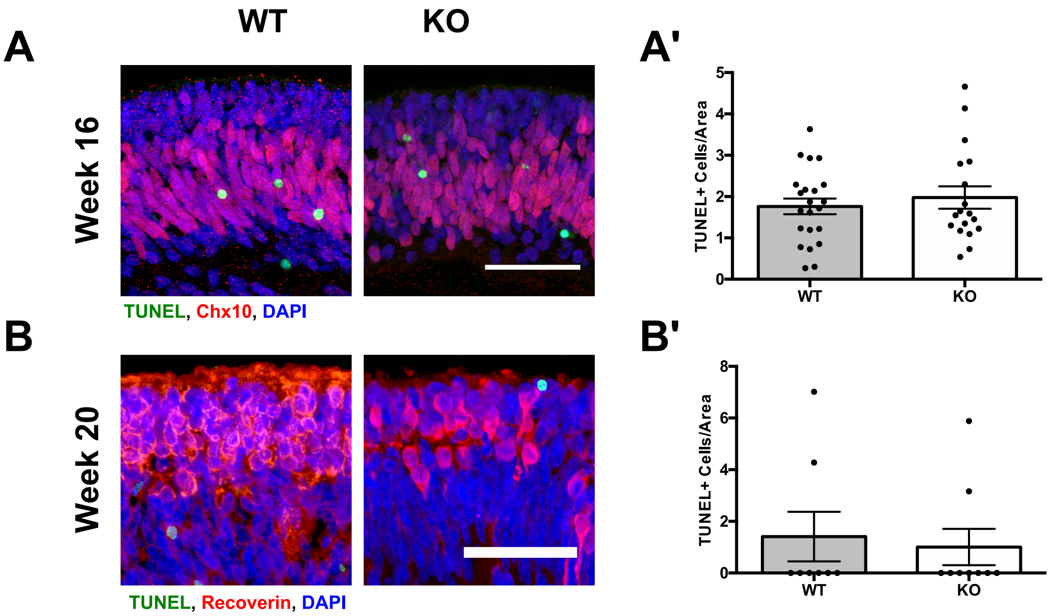

### Figure S7

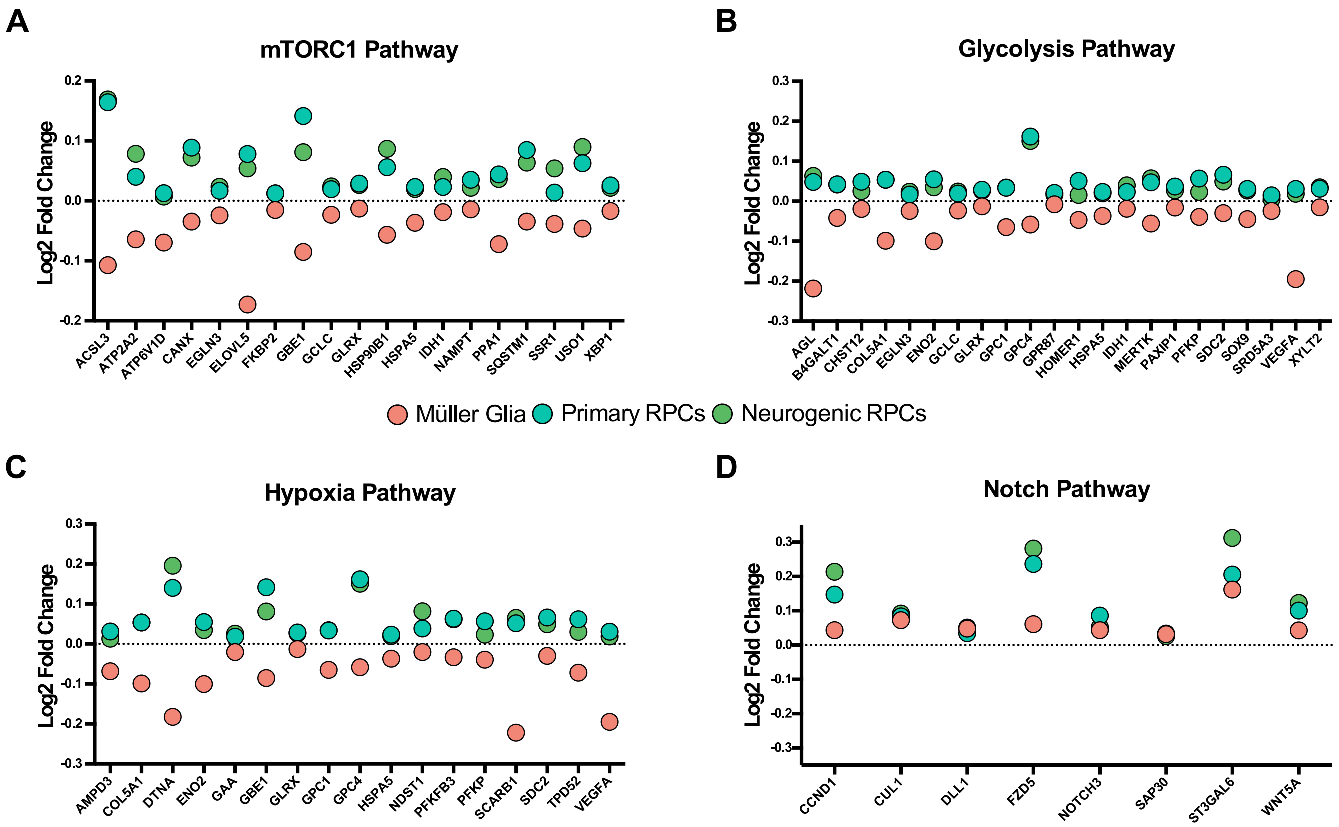
