## Supplementary material for "Multi-omic Analysis of Developing Human Retina and Organoids Reveals Cell-Specific Cis-Regulatory Elements and Mechanisms of Non-Coding Genetic Disease Risk": Figure S3

**A**

### Human and Organoid Candidate Disease-Gene-Associated CREs

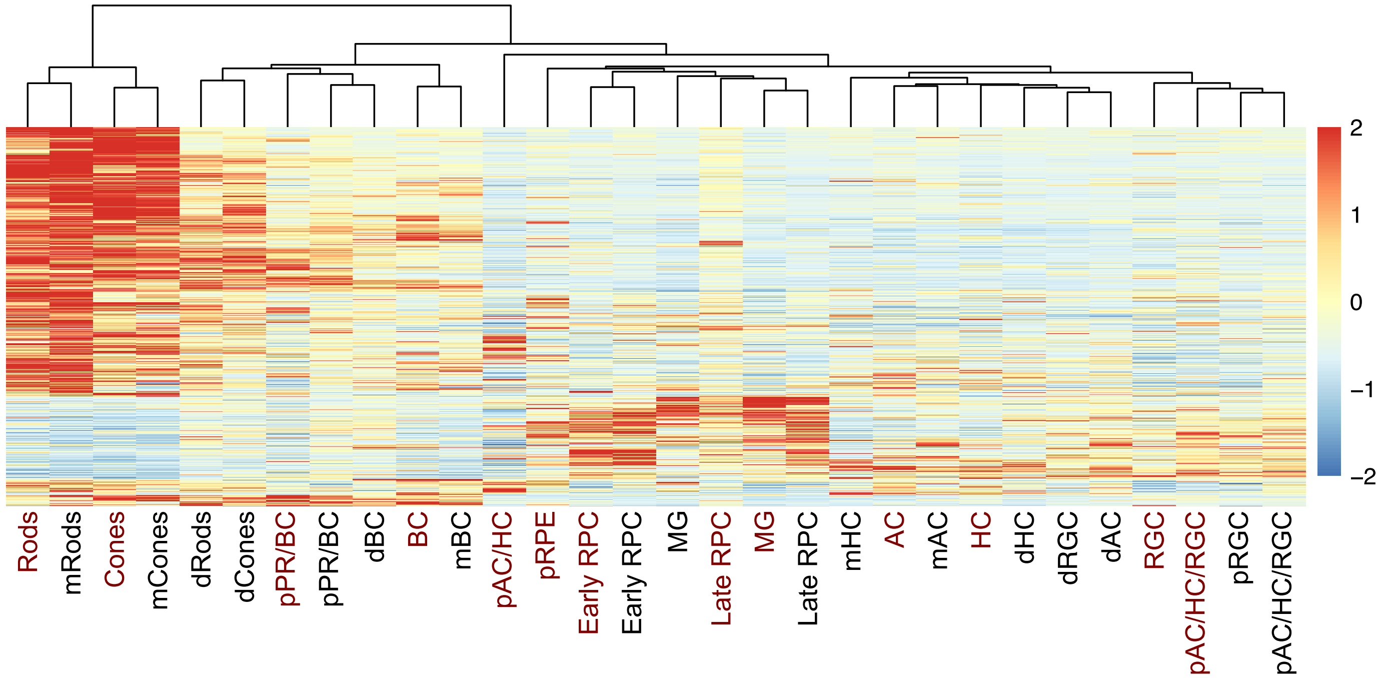


**B**

### Human and Mouse Candidate Disease-Gene-Associated CREs

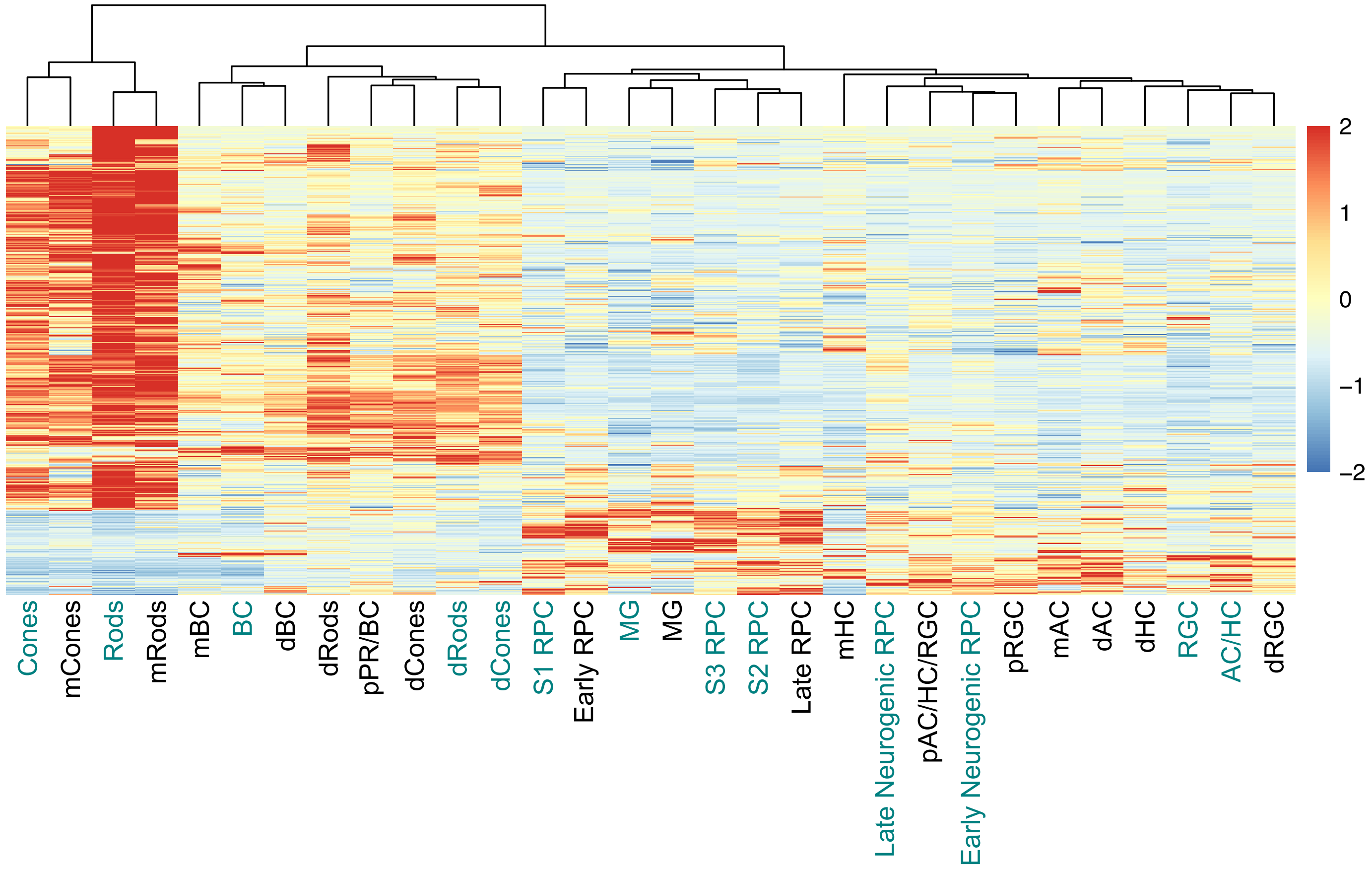
